# A Computational Model of Normal and Impaired Lexical Decision: Graded Semantic Effects

**DOI:** 10.1101/708156

**Authors:** Ya-Ning Chang, Steve Furber, Matthew Lambon Ralph, Stephen Welbourne

**Author notes:** Correspondence to: Ya-Ning Chang, MRC Cognition and Brain Sciences Unit, University of Cambridge, Cambridge CB2 7EF UK.

## Abstract

Lexical decision is an important paradigm in studies of visual word recognition yet the underlying mechanisms supporting the activity are not well understood. While most models of visual word recognition focus on orthographic processing as the primary locus of the lexical decision, a number of behavioural studies have suggested a flexible role for semantic processing regulated by the similarity of the nonword foil to real words. Here we developed a computational model that interactively combines visual-orthographic, phonological and semantic processing to perform lexical decisions. Importantly, the model was able to differentiate words from nonwords by dynamically integrating measures of polarity across the key processing layers. The model was more reliant on semantic information when nonword foils were pseudowords as opposed to consonant strings. Moreover, the model was able to capture a range of standard reading effects in lexical decision. Damage to the model also resulted in reading patterns observed in patients with pure alexia, phonological dyslexia, and semantic dementia, demonstrating for the first time that both normal and neurologically-impaired lexical decision can be addressed in a connectionist computational model of reading.

## Introduction

### 1.1. Lexical Decision

Lexical decision (LD) has been widely used to investigate the cognitive processes involved in visual word recognition. The task generally requires participants to make YES responses for words while NO responses for nonwords. Differences in measures of accuracy and response time are thought to illuminate the underlying processing of words. However, these measures can vary substantially depending on the lexicality of the nonword foils. For example, accurate and rapid responses can be observed when a word is tested in the context of consonant strings (i.e., orthographically illegal and unpronounceable, e.g., FJK), but these responses will become slower and less accurate when the same word is tested in the context of pseudowords (i.e., orthographically legal and pronounceable, e.g., FET), or of pseudohomophones (i.e., orthographically legal and sounding like real words, e.g., FEA) (Evans, Lambon Ralph, & Woollams, 2012; James, 1975; Ratcliff, Gomez, & McKoon, 2004; Shulman & Davison, 1977; Stone & Van Orden, 1993). These results suggest some control when making decisions in different contexts (Grainger & Jacobs, 1996; James, 1975; Plaut, 1997; Seidenberg & McClelland, 1989; Stone & Van Orden, 1993). For example, Stone and Van Orden (1993) reported that longer response latencies for words in the pseudohomphone condition compared to the pseudoword condition and then to the consonant string condition, demonstrating the influences of nonword lexicality. Further, the impact was larger for low than for high frequency words. It is likely that when judging words against consonant strings, the pathway from orthography to phonology is useful for indicating pronounceability. Whereas if nonwords are also pronounceable, the distinction cannot be made, and the pathway shift from orthography to lexical-semantics would be necessary. Thus, strategic control could be realised as the selection of reading pathways for making decisions (Stone & Van Orden, 1993).

Several studies have explored the influence of orthographic information on lexical decision (e.g., Andrews, 1989, 1992; Grainger & Jacobs, 1996). The most widely used orthographic measure is orthographic neighbourhood size (Coltheart, Davelaar, Jonasson, & Besner, 1977), defined as the number of words that can be created by replacing one constituent letter in a given word. Although there are different measures of orthographic structures (e.g., body neighbours and bigram/trigram frequencies) as potential confounders, it has been reported that decisions become difficult when nonwords have many orthographic neighbours, but easier when nonwords have few orthographic neighbours. This is particularly true when words are low frequency (see Andrews, 1997 for a review). However, the faciliatory effect may be reduced when accuracy rather than speed is emphasised (Binder et al., 2003).

There is also considerable evidence of phonological activation in visual lexical decision (Coltheart, Besner, Jonasson, & Davelaar, 1979; Meyer, Schvaneveldt, & Ruddy, 1974; Milota, Widau, McMickell, Juola, & Simpson, 1997; Patterson & Marcel, 1977; Rubenstein, Lewis, & Rubenstein, 1971). A relatively slow response is often observed for pseudohomophones compared with pseudowords, presumably because the lexical-semantic representations of real words are activated by the phonology of pseudohomophones (Grainger, Muneaux, Farioli, & Ziegler, 2005; Martensen, Dijkstra, & Maris, 2005).

While there is consistent evidence showing that visual orthography and phonology all play a role in lexical decision, the involvement of semantic processing in lexical decision has been somewhat debatable. It was originally thought that the retrieval of word meanings could only occur after lexical access has completed (Becker, 1980; Collins & Loftus, 1975; Forster, 1976; Morton, 1969). On this view, lexical access and meaning retrieval are strictly sequential and separate. However, numerous studies have investigated different aspects of semantic effects on lexical decision. For instance, imageability (or concreteness) effects (Balota et al., 2004; Cortese & Khanna, 2007; Degroot, 1989; Evans, Lambon Ralph, & Woollams, 2012; James, 1975; Kroll & Merves, 1986; Whaley, 1978), where highly imageable (or concrete) words (e.g., ball) are processed more quickly than abstract words (e.g., justice). Another example is semantic priming (Evans, Lambon Ralph, & Woollams, 2012; Joordens & Becker, 1997; Lupker & Pexman, 2010; Meyer & Schvaneveldt, 1971; Shulman & Davison, 1977; Yap, Tse, & Balota, 2009), where words with a semantically related prime are responded to faster relative to an unrelated prime. There are also studies investigating semantic ambiguity advantage (Azuma & Van Orden, 1997; Balota & Chumbley, 1984; Borowsky & Masson, 1996; Chumbley & Balota, 1984; Hino & Lupker, 1996; Hoffman & Woollams, 2015; Jastrzembski, 1981; Kellas, Ferraro, & Simpson, 1988; Millis & Bution, 1989; Rubenstein, Garfield, & Millikan, 1970). Furthermore, the extent of semantic involvement in lexical decision would seem to change under different contexts. The first study to explore graded semantic influences on lexical decision was conducted by James in 1975. He showed a reliable concreteness effect during lexical decision when using pseudoword and pseudohomophone foils, while the effect disappeared when testing with consonant strings. Some subsequent studies have replicated these findings using semantic priming under different foil conditions (Degroot, 1989; Joordens & Becker, 1997; Kroll & Merves, 1986; Shulman & Davison, 1977) while others have failed to find such effects (Lupker & Pexman, 2010; Yap, Tse, & Balota, 2009). More recently, Evans, Lambon Ralph and Woollams (2012) systematically investigated how imageability and semantic priming affect lexical decision in different nonword foil conditions. In their study, three different types of nonword foils were used including consonant strings, pseudowords, and pseudohomophones. Those foils were well controlled in their similarity to real words both orthographically and phonologically. Evans et al. (2012) demonstrated that semantic involvement in lexical decision was graded by the difficulty of the decisions under different conditions. There were stronger semantic effects with pseudohomophones than with pseudowords, and the effects were stronger with pseudowords than with consonant strings. These results confirmed semantic influences on lexical decision in circumstances where the nonwords are plausible. The authors suggested that the discrepant reports of semantic effects in the literature might be because the nonword stimuli have rather different ranges of orthographic neighbourhood and/or bigram frequency across studies and the effects seem to be more pronounced when the nonwords have higher orthographic neighbourhood size and/or bigram frequency.

There is also evidence of semantic involvement in lexical decision from neuroimaging studies. Binder et al. (2003) found stronger activation for words than nonwords in the dorsal prefrontal cortex, the inferior temporal and posterior middle temporal gyri – all areas previously associated with semantic processing (Binder, Desai, Graves, & Conant, 2009; Binder et al., 1999; Vandenberghe, Price, Wise, Josephs, & Frackowiak, 1996). Similarly, Woollams, Silani, Okada, Patterson and Price (2011) detected left anterior temporal activation for orthographically atypical words relative to orthographically typical words when lexical decisions were made more difficult in the context of pseudohomophone foils, in that orthographical typicality was defined in terms of summed bigram frequency. This left anterior temporal region has been identified as a semantic “hub” combining various sensory and motor information to form transmodal semantic representations (Patterson, Nestor, & Rogers, 2007; Rogers, Lambon Ralph, Hodges, & Patterson, 2004; Visser, Jefferies, & Lambon Ralph, 2010). A similar effect has also been detected in EEG where atypical words were found to elicit stronger source currents than typical words at 160 ms with the effect being driven by sources in the left inferior anterior temporal lobe (Hauk et al., 2006). The main effect of lexicality was observed at a later time frame around 200 ms, and it was further confirmed by their subsequent EEG/MEG study (Hauk, Coutout, Holden, & Chen, 2012). In addition, Fujimaki and colleagues (2009) used an fMRI assisted MEG technique to explore differences in activations elicited by a phonological decision task (detecting vowels in nonwords) and a lexical decision task. Significantly different brain activations were found in the left anterior temporal area between 200-400 ms.

These effects are consistent with observations of patients with semantic dementia (SD), who have bilateral atrophy of the anterior temporal lobes. These patients show a progressive degeneration of semantic knowledge (Hodges, Patterson, Oxbury, & Funnell, 1992). When patients are asked to perform two-alternative forced-choice visual lexical decision, they can make a decision correctly when a word has a higher bigram/trigram frequency than its nonword pair but have difficulty in the reverse condition (Patterson et al., 2006; Rogers, Lambon Ralph, Hodges, et al., 2004). Overall, these results suggest semantic information is used in lexical decision, but to a varying extent depending on the exact nature of the decision-making difficulties. Semantic processing is particularly likely to be engaged to assist recognition of atypical words, or for the rejection of word-like nonwords. The issue as to how semantics interacts with other processes arising from the underlying machinery in the recognition system for lexical decision remains to be systematically investigated.

### 1.2. Computational Models of Lexical Decision

Several models of lexical decision have been proposed, based on theories of visual word recognition (Coltheart, Davelaar, Jonasson, & Besner, 1977; Coltheart, Rastle, Perry, Langdon, & Ziegler, 2001; Grainger & Jacobs, 1996; Perry, Ziegler, & Zorzi, 2007; Plaut, 1997; Seidenberg & McClelland, 1989). These can be broadly divided into two groups depending on whether they assume the decision is made using some kind of lexical lookup procedure, or whether it is made based on more distributed properties of the underlying representations.

#### 1.2.1. Models Based on Lexical Accounts

Many researchers argue that lexical decision is made by checking whether a letter string has a corresponding lexical entity in the orthographic lexicon (Coltheart, et al., 1977). On this view, activation within the orthographic lexicon is the key factor in lexical decision. Phonology may also be involved, but it is a relatively late process after the mental lexicon search (Coltheart, Besner, Jonasson, & Davelaar, 1979), while the semantic system is generally not involved in these models (Coltheart, et al., 2001). This view is shared by Grainger and Jacobs (1996), who developed a computational model of lexical decision based on the interactive activation (IA) model (McClelland & Rumelhart, 1981). The word layer in the IA model could be considered as an orthographic lexicon where each input word form has a corresponding word unit. Grainger and Jacobs’s (1996) multiple read-out model (MROM) extended the IA model to perform the lexical decision task by adopting three activation criteria to determine the type of stimulus (i.e., word/nonword) and speed of a response. A word response could be made either when the particular word unit activation reached a local criterion, M, or the overall activity in the word layer reached a global criterion, Σ, before the temporal deadline T. The RT was based on the earliest moment where either of criteria was met. Following the IA model, the resting-level activations of word units varied as a function of word frequency. In addition, Grainger and Jacobs (1996) assumed that the local criterion M should be fixed, while the global criterion Σ and the temporal deadline T would vary according to the lexical frequency status of the stimulus. MROM was able to simulate several standard effects seen in lexical decision including frequency effects, orthographic neighbourhood size effects, and their interactions (Grainger and Jacobs, 1996). Other models of visual word recognition such as the dual-route cascaded (DRC) model (Coltheart, et al., 2001) and the connectionist dual process (CDP+) model (Perry, Ziegler, & Zorzi, 2007) also share the similar decision mechanisms to that in the MROM model. However, one major problem with the MROM model and its variants is related to the use of the temporal deadline mechanism, which inevitably means that the models cannot generate variable nonword decision times when testing different types of nonword stimuli (Ratcliff, Gomez & Mckoon, 2004; Wagenmakers et al., 2008).

Note that, within this framework, some researchers (Blazely, Coltheart, & Casey, 2005; Borowsky & Besner, 1993) have proposed that the semantic system can be involved in lexical decision and is responsible for the effects of semantic priming and imageability in lexical decision (e.g., James, 1975; Meyer, Schvaneveldt, & Ruddy, 1975). The decision could be made either by monitoring the activation within the semantic system (Borowsky & Besner, 1993) or via the feedback to the orthographic lexicon (Blazely, Coltheart, & Casey, 2005). However, this has yet been implemented to any localist model, so it remains unclear as to actual mechanisms that underpin the decisions based on the semantic system and its interaction with the orthographic lexicon.

#### 1.2.2. Models Based on Distributed Views

An alternative theory of visual word recognition argues that there is no mental lexicon for the store of word knowledge in the recognition system (Dilkina, McClelland, & Plaut, 2010; Plaut, 1997; Rogers, Lambon Ralph, Hodges, et al., 2004; Seidenberg & McClelland, 1989, 1990). On this view, the fundamental differences between words and nonwords are the nature of their underlying representations. The lexical decision can be made on the basis of the differential activations elicited by familiar words and unfamiliar nonwords. When presenting a word, strong activations are expected, because the mappings between the visual or orthographic representation of the word and its phonological and semantic representations have been learned. On the other hand, relatively weaker activations would be found for novel nonword representations. Both words and nonwords will activate visual, orthographic, phonological and semantic representations and these different levels of representations interact with each other during lexical decision. A critical difference between this view and the lexical view is that semantic processing is considered important for lexical decision (Dilkina, et al., 2010; Rogers, Lambon Ralph, Hodges, et al., 2004) in addition to orthographic and phonological processing (Plaut, 1997; Seidenberg & McClelland, 1989). Within this framework, several computational models have been developed to simulate the processes of lexical decision (Dilkina, et al., 2010; Harm & Seidenberg, 2004; Plaut, 1997; Seidenberg & McClelland, 1989). Plaut (1997) proposed that the measure of how strongly units were activated, called stress or polarity, could be used as a basis for making lexical decisions. Plaut developed a feedforward model, which consisted of orthographic, phonological and semantic components and demonstrated that words tended to produce higher stress than nonwords at the semantic layer allowing a discrimination rate of over 95%. In addition, the network tended to produce higher semantic stress for pseudohomophones than for pseudowords. This might explain why pseudohomophones are more difficult to reject (Coltheart, et al., 1977; Meyer, et al., 1974; Milota, et al., 1997; Patterson & Marcel, 1977; Rubenstein, et al., 1971). Subsequently, Harm and Seidenberg (2004) developed a fully implemented reading model to explore how the skilled readers generate meanings of words from print by incorporating a pre-existing knowledge of the interactive mappings between phonology and semantics with newly learned mappings from orthography to phonology and orthography to semantics. Although lexical decision was not the main focus in their paper, they explored how the model could account for the pseudohomophone effect in lexical decision. They proposed that lexical decision could be made by determining the discrepancy between the input orthographic patterns and the orthographic patterns recreated from semantics. If the feedback information was consistent with the orthographic information, a word response was made; otherwise, a negative response was given. Dilkina, McClelland and Plaut (2010) also developed a connectionist single-system model of semantic and lexical processing to simulate lexical decision performance in SD patients. The model included four input/output layers including vision, orthography, phonology and action layers. There were also two hidden layers, one integrative semantic layer and an intermediate layer that facilitates the mapping between orthography and phonology. The lexical decisions were made in a similar way to Harm and Seidenberg (2004) by measuring the orthographic feedback, termed orthographic echo, for the presented words and nonwords. It is worth noting that although this measure of lexical decision is recorded at the orthographic layer, the orthographic echo is also dependent on activations at the semantic layer.

Several researchers have challenged the distributed view of lexical decision by arguing that these models would not be able to account for some patients with semantic impairments who show normal lexical decision accuracy (Coltheart 2004; Blazely, Coltheart, & Casey, 2005; Borowsky & Besner, 2006). That is if the semantic system is important for lexical decision as advocated by the distributed view, when the system is damaged the lexical decision performance should be greatly affected. To address this issue, Plaut and Booth (2006) developed a model that involved mappings from orthography to semantics. They applied a range of damage to the model’s semantic layer. The results showed that the models’ lexical decision performance was only slightly decreased with increasing semantic damage, whereas the model’s semantic performance was sustainably deteriorated. As explained by Plaut and Booth, the model with semantic impairment was still able to distinguish words from nonwords because of the much stronger activations of words compared to that of nonwords, though the activations between words were not easily distinguishable.

#### 1.2.3. Accumulated Information for Lexical Decision

Other models have emphasised the use of accumulated information for decision-making tasks (Busemeyer & Townsend, 1993; Norris, 2006, 2009; Ratcliff, Gomez and Mckoon, 2004; Usher & McClelland, 2001, 2004). In some accumulator models, decisions are made by diffusion processes that continuously accumulate information over time from a starting point toward response boundaries. It can be implemented with a simple diffusion process such as the diffusion model (Ratcliff et al., 2004), where positive evidence for one of the alternative responses can be considered negative evidence for the other alternative. Other accumulator models with two or more diffusion processes allow the evidence in the accumulator to decrease due to random noise (Busemeyer & Townsend, 1993) or inhibition between the processes (Usher & McClelland, 2001, 2004). Among these accumulator models, the diffusion model (Ratcliff et al., 2004) has been widely applied to account for behavioural data in lexical decision. To choose between words and nonwords, the speed of information accumulation in the diffusion model, called the drift rate, is determined by the lexical status of the stimuli. The more word-like, the stimulus the higher the drift rate. The diffusion model was able to account for a range of phenomena seen in lexical decision including the word frequency effects and the correct patterns of RT distributions for both words and various types of nonwords.

Another type of accumulator model is implemented on the basis of Bayesian probability, named the Bayesian reader (Norris, 2006, 2009). It assumes that subjects would consistently compute the probability of the stimulus being a word or a nonword based on its lexical status. The word likelihood is the sum of the probabilities of all possible letter strings whereas the nonword likelihood is simply one minus word probability (Norris, 2009). The Bayesian reader performed reasonably well in lexical decision and was able to simulate the effects of word frequency and the nature of the nonwords reported in Ratcliff et al.’s (2004) experiments. However, the model was unable to simulate the distribution of RTs for incorrect lexical decisions. It is also unclear how the model would perform when it is tested against nonwords with high orthographic neighbourhood sizes.

### 1.3. Summary

Evidence from behavioural, neuroimaging and patient studies, suggest that orthographic processing, phonological processing and lexical-semantic processing are all involved in lexical decision albeit to different extents dependent on the properties of word stimuli and the foil types. Human readers are able to flexibly use the available information from the recognition processes to support efficient lexical decision. Readers may rely on orthographic information if it is reliable and sufficient for decisions; otherwise, they may need to utilise phonological and semantic information (Plaut, 1997; Seidenberg & McClelland, 1989). While previous models of lexical decision have demonstrated how lexical decision can be made based on output generated from a single processing component, no model has yet shown how information from all the processing components of word recognition can be flexibly combined to support lexical decisions. This is the first time that these ideas have been implemented computationally. We aimed to develop a fully implemented computational model of visual word recognition that interactively combines visual, orthographic, phonological and semantic processing based on the parallel distributed processing (PDP) framework (Chang, Furber, & Welbourne, 2012a; Welbourne & Lambon Ralph, 2007; Plaut, McClelland, Seidenberg, & Patterson, 1996; Seidenberg & McClelland, 1989) to account for a range of standard effects in visual lexical decision tasks including frequency, consistency, orthographic neighbourhood size, and word length (number of letters) (Andrews, 1982; Balota et al. 2004; ; Cortese & Khanna, 2007; Seidenberg, et al., 1984; Waters & Seidenberg, 1985). In particular, we sought to use the model to investigate the graded semantic effects in the contexts of different foil types as reported by Evans et al. (2012). We also tested if damage to visual-orthographic, phonological and semantic layers in the model could result in similar impaired lexical decision performance as that observed in patients with corresponding functional impairments, which are pure alexia, phonological dyslexia and semantic dementia.

Two simulations were run to investigate the models’ performance in the following areas: (1) Normal lexical decision – including effects of frequency, consistency, foil type, imageability, word length and orthographic neighbourhood size; (2) Impaired lexical decision – including a comparison of how damage to the different processing areas could be mapped onto the lexical decision performance of different patient groups. An additional simulation reported in Appendix A was conducted to investigate the role of control units in regulating the temporal dynamics of the model. All of the simulations used the same network architecture (Figure 1), which is fully described in Simulation 1.

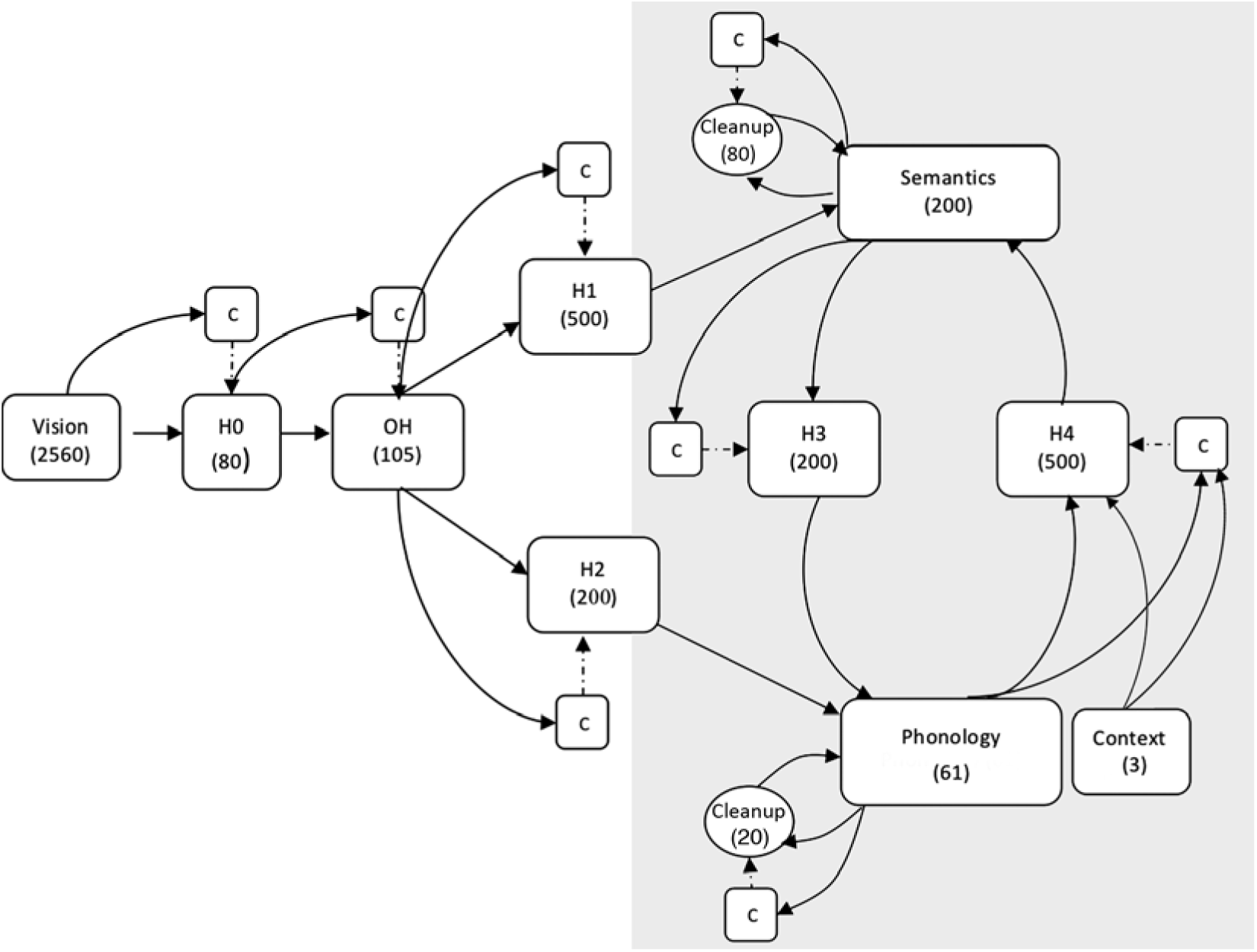

## 2. Simulation 1

Simulation 1 was designed to develop a recurrent model of word reading based on the general triangle framework (Harm & Seidenberg, 2004; Plaut et al., 1996; Seidenberg & McClelland, 1989; Welbourne & Lambon Ralph, 2007; Welbourne, Woollams, Crisp, & Lambon Ralph, 2011), but crucially including a visual level of representation similar to that used by Chang et al (2012a). In that study the addition of a visual layer of processing allowed the model to capture effects related to word length (measured by the number of letters), which have previously been problematic for this type of model to address, in particular, the length by lexicality effect (Weekes, 1997). However, that model did not include an interactive semantic processing layer; this is the first time that a full triangle model including visual, orthographic, phonological and semantic processing layers has been reported.

### 2.1. Method

#### 2.1.1. Network architecture

The model was a continuous recurrent network. The architecture of the model is shown in Figure 1. The model had two separate pathways for processing words from visual input: a phonological pathway and a semantic pathway. The word stimuli were presented to the visual layer of the model that connects to the OH layer via a set of 80 hidden units. The OH layer used here is equivalent to the orthographic layer in the triangle model, except that the orthographic representations were learned through the course of training (as in human development), rather than being supplied as inputs. It connects to both phonological and semantic units via two more sets of hidden units. Both the phonological and semantic units were connected to their own set of clean up units, which allowed for the development of phonological and semantic attractors in the model. In addition, there were three context units, which were used to provide contextual information for disambiguating homophones: While there is no way to distinguish meanings of homophones in single word reading, during natural reading the context will almost always give sufficient cues. Phonological and semantic units were connected to each other via two hidden unit layers. All output layers in the model were given a fixed negative bias of −2 to encourage sparse representations. Perhaps the most unusual element of the architecture is the control units associated with each layer except input and output layers. These units receive the same inputs as the layer they are connected to, and all their outgoing connections are inhibitory, allowing them to control the activation of all the units in a layer simultaneously. These units turn out to be critically important for training the present recurrent network as they help to regularise the time dynamics in the network by preventing activations in the deep layer from causing unwanted disturbance to the processing in the earlier layers. The details are illustrated in Appendix A.

In the current model, the activation values of all units were updated concurrently and ramped up or down gradually over time. The dynamics of the network was implemented by using an output integrator equation (Pineda, 1987). The output integrator function can be written as follows:

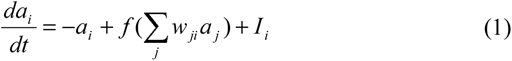

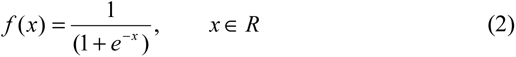

where *a_i_* is the current output of the unit *i*, where 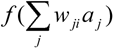 is the summed output contribution from other units *j*, where *f* is a logistic function, and where *I_i_* is a constant and represents an external input bias. For computational simulations, the continuous time is generally approximated by using discrete time steps, in that some intervals of the processing time are sampled and broken into a number of finite ticks. In the current reading simulation, the network was run for ten intervals of time, and three ticks per interval were used for the approximation of the continuous time in the network.

#### 2.1.2. Visual representations

The training corpus consisted of 2,971 monosyllabic words. The visual representations used here were adapted from those used in Chang et al.’s study (Chang et al., 2012a). The network was directly fed with bitmap images of words in Arial 12-point lower case font, represented in white against a black background. Each word was positioned with its vowel aligned on the central slot of the image^1^. For words that have a second vowel (e.g. boat), the second vowel was placed right next to the first vowel. Ten 16×16 pixel slots were used so there were 2,560 visual units.

#### 2.1.3. Phonological representations

The scheme of phonological representations was the same as that used in the Plaut et al. (1996) model. Each word was parsed into onset, vowel and coda clusters of phonemes with specific units used to represent each possible phoneme in each cluster (Table 1). This gives a total of 61 phoneme units.

**Table 1.** Orthographic and Phonological Representations

|  |  |
| --- | --- |
| Orthography |  |
| Onset | Y S P T K Q C B D G F V J Z L M N R W H CH GH GN PH PS RH SH TH<br>TS WH |
| Vowel | E I O U A Y AI AU AW AY EA EE EI EU EW EY IE OA OE OI OO OU OW<br>OY UE UI UY |
| Coda | H R L M N B D G C X F V J S Z P T K Q BB CH CK DD DG FF GG GH<br>GN KS LL NG NN PH PP PS RR SH SL SS TCH TH TS TT ZZ U E ES ED |
| Phonology |  |
| Onset | s S C z Z j f v T D p b t d k g m n h l r w y |
| Vowel | a e i o u @ ^ A E I O U W Y |
| Coda | r l m n N b g d ps ks ts s z f v p k t S Z T<br>D C j |

#### 2.1.4. Context representations

The three context units were used to help differentiate the meanings of homophones. For nonhomophones, the context units were all set to zero. Within the same homophone family, different context units were assigned, one for each. The maximum number of meanings corresponding to a given pronunciation in the training corpus is four.

#### 2.1.5. Semantic representations

We utilised a set of representations previously developed by Chang, Furber, and Welbourne (2012b). These were developed by searching for a representational structure that met five key criteria that were considered desirable for use in large scale computational models: binary coding; sparse coding; a fixed number of critical semantic features in each vector; scalable vectors and most importantly preservation of the human-like semantic structure.

In the literature, several semantic representation schemes have been proposed either based on feature norms (Garrard, Lambon Ralph, Hodges, & Patterson, 2001; McRae, Cree, Seidenberg, & McNorgan, 2005), statistical co-occurrence semantic analysis (Landauer, Foltz, & Laham, 1998; Lund & Burgess, 1996; Rohde, Gonnerman, & Plaut, 2006), artificially generated patterns (Dilkina, et al., 2010; Plaut, 1997; Rogers, Lambon Ralph, & Garrard et al. 2004; Welbourne, et al. 2011) or Miller, Beckwith, Fellbaum, Gross, and Miller’s (1990) online WordNet database (Harm and Seidenberg, 2004).

Chang et al. (2012b) considered that only schemes that derived latent semantic relations based on co-occurrence matrices from text corpora could meet all the desirable criteria and that the most promising candidate among these was Correlated Occurrence Analogue to Lexical Semantics (COALS; Rohde et al., 2006). COALS uses singular value decomposition (SVD) on co-occurrence statistics to produce an arbitrary length semantic vector for each word. In the original formulation, only the positive valued vectors were used. This means that the information contained in negative parts of the vector could be lost. The fact that *dogs* have four legs would be captured, but the fact that they never fly would not. It is because *dogs* are more likely to co-occur with legs but not with fly. Chang et al. explored the possibility that including the negative features might improve the quality of the semantic representations. They started with a 100-dimensional vector from COALS, which they duplicated to create a 200-dimensional vector. The first 100-dimensions coded the top *n* positive features and the second 100-dimensions coded the top *m* negative features. The optimum number of positive and negative components was assessed by comparing the semantic categories within the artificial semantic codes with six groups of items categorised by human subjects in the Garrard et al.’s (2001) study.

Coding schemes that involved negative items consistently produced category structures that more closely matched the human results. The set of semantic vectors that most closely matched the human-derived structure was found to use five positive and fifteen negative features. Hierarchical clustering analysis showed that compared with Garrard et al. data, most items were clustered into the same group as in the human category. For example, living thing items were separated from nonliving thing items. Within the living thing items, the items (e.g. *dog*) in the animal group and the items (e.g. *apple*) in the fruit group were well separated. Within the nonliving things, the broader category of tools (e.g. *hammer*) was well distinguished from the group of vehicles (e.g. *train*), although the boundary between tools and household items was less clear. More details can be found in Chang et al.’s (2012b) paper.

The semantic representations used in the present simulations were generated using the scheme described above. The meaning of each word was represented by a 200-dimensional semantic vector. Each vector had five active units in the first half of the vector converted from the top five positive attributes and fifteen active units in the second half of the vector converted from the top fifteen negative attributes.

#### 2.1.2. Training procedure

The training was separated into two phases. In phase 1, the links between phonology and semantics were trained (shown in grey in Figure 1). This phase of training was intended to correspond to pre-literate language learning in children. In phase 2 the full reading model was trained.

In phase 1, the phonology-semantics model was subdivided into two parts: the production model learning the mappings from semantics to phonology, and the comprehension model learning the mappings from phonology to semantics (Figure 2). Both parts were trained on the full corpus of 2,971 monosyllabic words. The probability of each word being presented to the model was determined by its logarithmic frequency. Slightly different learning rates and weight decays were used to train the two parts: the production model was trained with a learning rate of 0.2 and a weight decay of 1E-7, while the comprehension model trained with a learning rate of .05 and weight decay of zero. The initial weights were randomly set to values between −0.1 and 0.1.

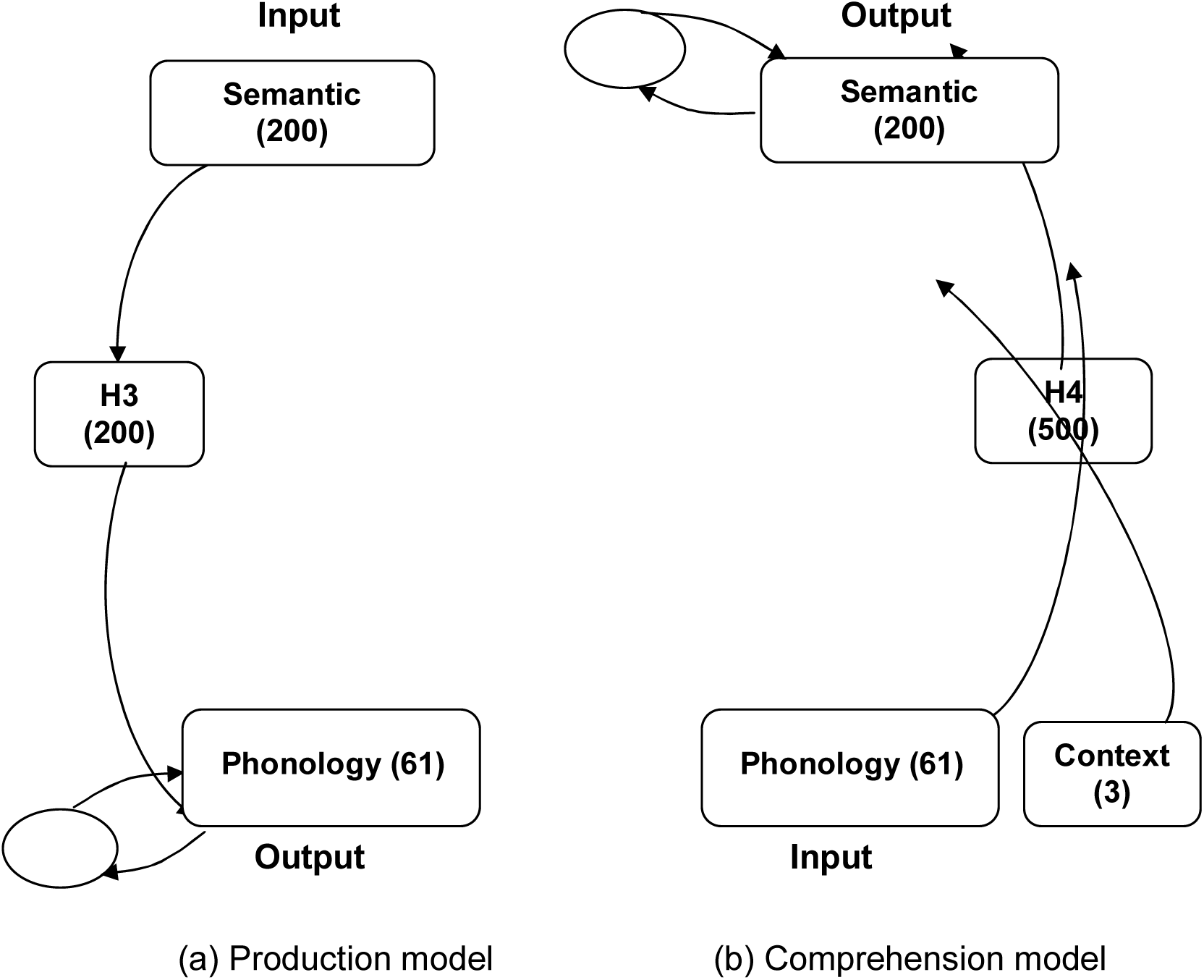

Each example was presented for six intervals of time and each interval of time was divided into three ticks. In each presentation, the input pattern was clamped onto the appropriate units for six intervals of time. For the last two intervals, the activations of output units were compared with their targets. Error score was measured on the basis of divergence (cross-entropy; Plaut et al. 1996) between the target and the actual activation of the output units. It was used to calculate weight changes according to the back-propagation through time (BPTT) algorithm (Pearlmutter, 1989, 1995). No error was recorded if the output unit’s activation and target were within 0.1 of each other. After this training, additional 40,000 epochs of interleaved training on the mappings between phonology and semantics was applied in order to fine-tune the combined phonology-semantics model.

In phase 2, the full reading model was trained using the BPTT algorithm with a learning rate of 0.1, a weight decay of 1E-8 and a momentum of 0.9. The visual representation of a word was presented at the input units for ten intervals (again each interval of time was broken into three ticks). The task was to produce correct phonological and semantic patterns. For the last two intervals, the output units were compared with their corresponding phonological, or semantic targets and errors were computed. No error was computed when the output unit’s activation and target were within 0.001. Again logarithmic frequency was used to determine the probability with which word was presented to the model. To preclude any possibility that simulation result could be generated from one particular set of initial weights, twenty models with different initial weights were trained. When the improvement of the model’s performance on phonology and semantics had slowed down and had reached an asymptote, following Bishop (2006), the accuracy rate on regular nonword pronunciations was used to determine the end point of training.

#### 2.1.3. Testing Procedures

The testing procedures for both training phases were precisely the same. The decoding procedure for semantics was based on the Euclidean distances between the activations of the semantic units and each of the semantic representations in the training corpus (Monaghan, Shillcock, & McDonald, 2004; Monaghan et al., 2017). The semantic representation which was closest to the activation of the semantic units was taken as the semantic output. If the output was the same as the target representation, it was a correct response. The procedure for the generation of the phonological output was the same as that used in Plaut et al.’s (1996) study. For the vowel units, the most activated vowel unit was selected as the output. Onset and coda units were divided into groups of mutually exclusive units and the highest active unit above 0.5 was taken as the output for each group. If no unit was active above 0.5 then the group did not contribute to the output. Finally, if either of the ks ts or ps unit was active along with their components, then the order of the components was reversed.

### 2.2. Training performance

After two million presentations, phase 1 training was halted. The accuracy rates of the production and comprehension model were 99.97% and 99.43% respectively. Figure 3 shows the performance of the full word reading model throughout phase 2 training with and without control units. For the model trained with control units, both semantic and phonological units learned relatively quickly (phonology was faster than semantics), and reached near-perfect performance after 600,000 epochs. The average accuracy rates for the model to produce correct phonological and semantic patterns in the word reading task were 99.3% and 97.4% respectively. By contrast, when the model was trained without using control units, the phonological units never achieved an accuracy rate higher than 91%, and the semantic units were still at only 20% accuracy after one million epochs. As we shall see in Appendix A, this is because without control units the system cannot make the most efficient use of the embedded knowledge in the pre-trained connections between semantics and phonology.

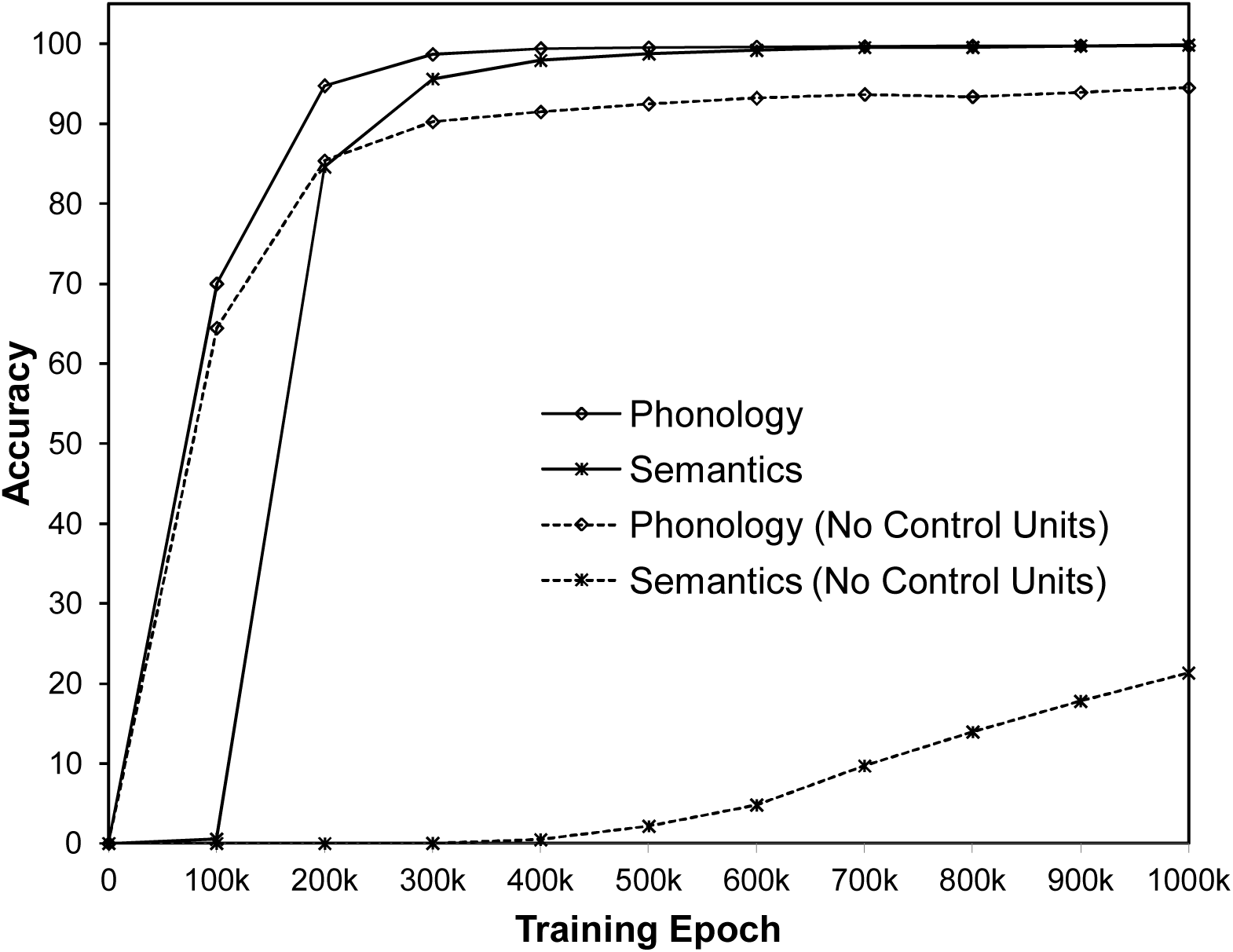

### 2.3. Exploring Polarity Measure for Lexical Decision

Plaut (1997) demonstrated that the parallel distributed models could perform the lexical decision task based on the measure of polarity in the semantic layer, which is essentially a test of how binary the representations are. The idea behind this is that during training, the units are trained to represent the target patterns consisting of binary values. Thus when unfamiliar items such as nonwords are presented, the units tend to remain closer to their initial states. To capture this phenomenon, Plaut (1997) used a formula to compute the index of unit binarisation, termed unit polarity:

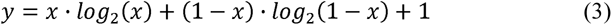

where *x* is the unit activation ranging from 0 to 1 and *y* is the polarity measure.

However, Plaut’s (1997) model only looked at polarity scores in the semantic layer, and it remains unclear if polarity scores based on visual, orthographic or phonological processing in the reading system would also be useful. Thus, in the present study, polarity scores across units in the H0 and OH (orthographic), phonological and semantic layers were integrated.

#### 2.3.1. Testing stimuli

To investigate if the individual polarity scores from different key processing layers and their combined scores could provide a reliable source for making lexical decisions, we computed the polarity values generated from a set of words and nonwords taken from Evans et al.’s (2012) study. They used three different types of nonword foils where the foils were controlled in their orthographic and phonological relationships to the real words. That allowed us to see the polarity differences between words and different sets of nonword foils. In Evans et al.’s study, there were 80 words. Each word had its matched consonant string, pseudoword and pseudohomophone. However, ten of these words were not in our training corpus, so these were removed from the test along with their matched nonword foils leaving a total of 70 words along with three sets of matched nonword foils (consonant strings, pseudowords and pseudohomophones).

#### 2.3.2. Average polarities at different layers

Figure 4 shows the average polarities as a function of time in each of the processing layers. For the orthographic layer (Figure 4, panel a), there were small differences between words and pseudowords or pseudohomophones; however, the differences between consonant strings and all other types of stimuli were relatively large, indicating that the distinction between words and consonant strings may be easier than that of words and pseudowords or pseudohomophones. The polarity scores in the phonological layer (Figure 4, panel b) were slightly larger for words and the smallest for consonant strings with pseudowords and pseudohomophones somewhere in between. All the differences were small. In the semantic layer (Figure 4 panel c), the main feature was that the polarity for words was higher than for all other stimuli, suggesting that this layer might make the most reliable contribution to lexical decision. However, it should be noted that these differences emerge relatively late compared with the difference between words and consonant strings in the orthographic layer. Overall these results suggest that different areas of the model may contribute differently to the lexical decision task, depending on whether it could provide sufficient information to be used for deciding which foils can be rejected, and when the decision occurs. Consonant strings can be distinguished from words early on in the visual-orthographic processing areas, whereas pseudowords and pseudohomophones may require input from semantic processing areas. By averaging the polarities across these processing layers, it might be possible to maximise the differences.

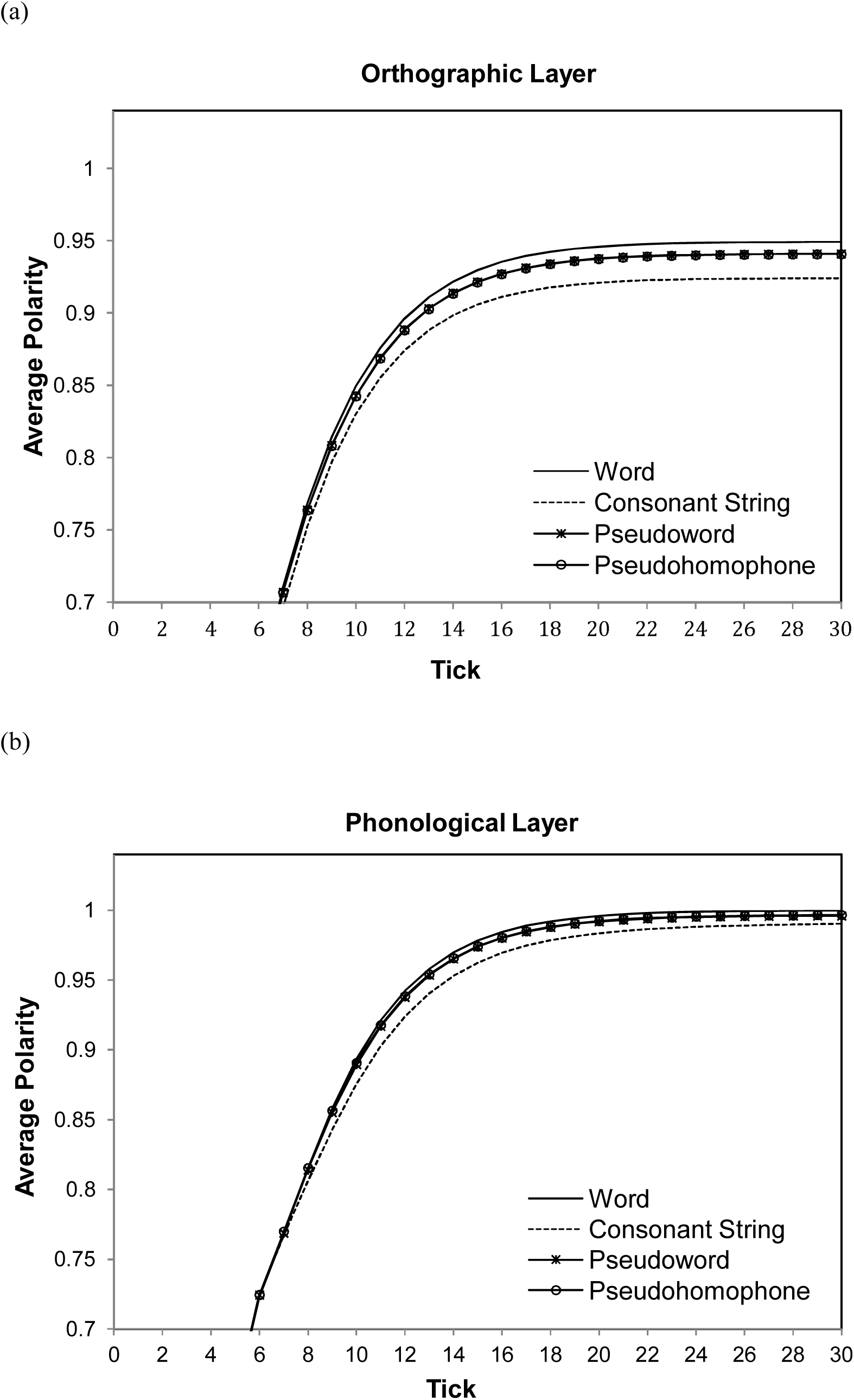
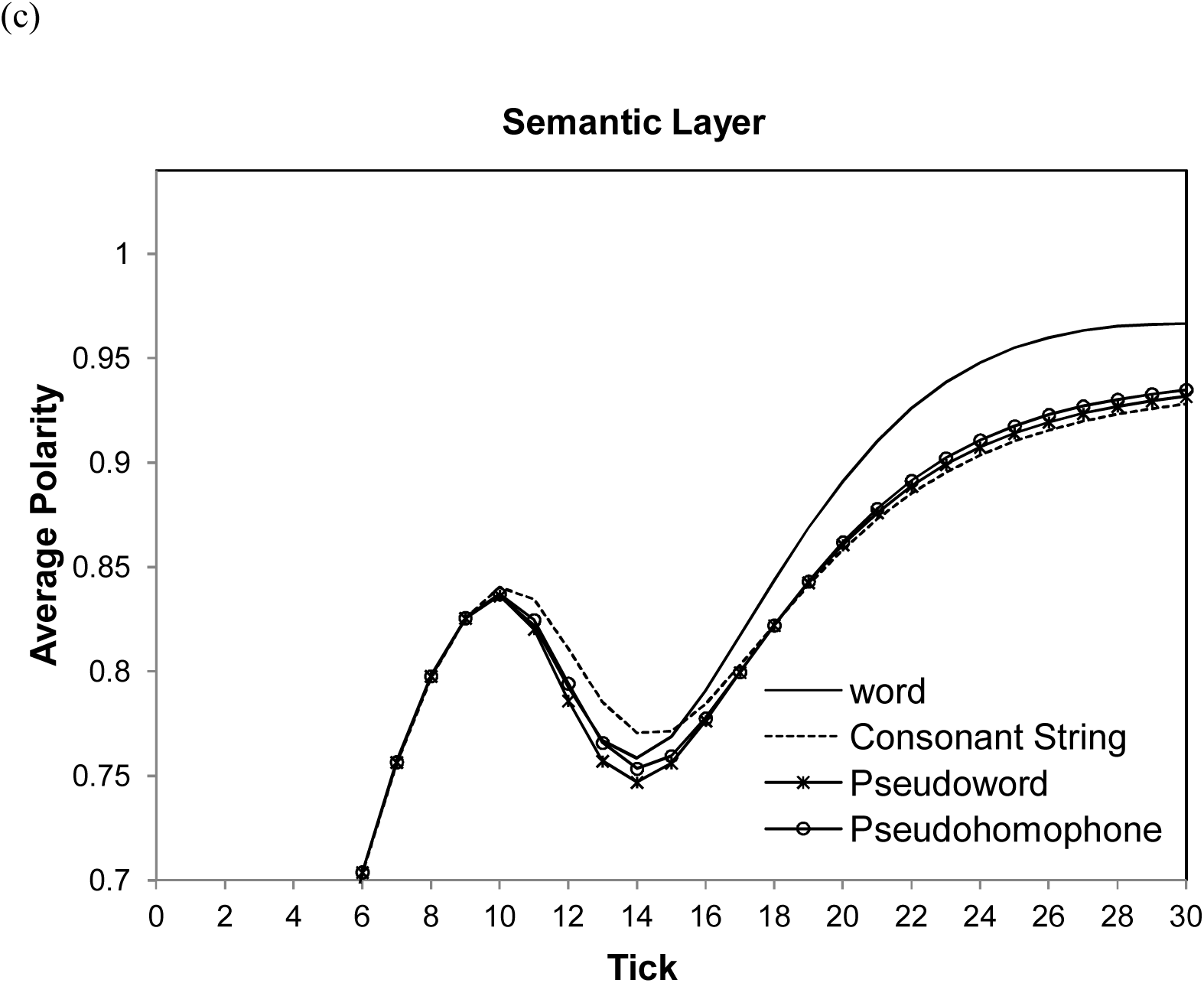

To test this possibility, average polarity differences were computed for the last two intervals of network time (i.e., from 25^th^ tick to 30^th^ tick). The difference was computed according to the following formula, which calculates the z score of the differences separately for each set of foils:

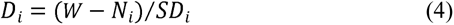

*i* refers the set of nonword foils (consonant string / pseudoword / pseudohomophone); *W* is the average word polarity across all items; *N_i_* is the average nonword polarity across all items in the set; *SD_i_* is the standard deviation of the nonword polarity for the items in that set.

Figure 5 shows the average polarity differences of the key processing layers by nonword condition. One-sample t tests were conducted to examine whether the average polarity differences in each condition was greater than zero. The results showed that all the differences were significantly different from zero (all *ps* < .05). A one-way repeated measures ANOVA analysis with nonword condition as a within-subject factor revealed that the effect of nonword condition was also significant, *F*(2, 38) = 163.44, *p* < .001. A Tukey post-hoc test revealed that the polarity difference in the consonant string condition (1.33) was significantly larger than that in both the pseudoword condition (0.90), *p* < .001, and the pseudohomophone condition (0.82), *p* < .001. Moreover, the polarity difference in the pseudoword condition was also significantly larger than that in the pseudohomophone condition, *p* < .05.

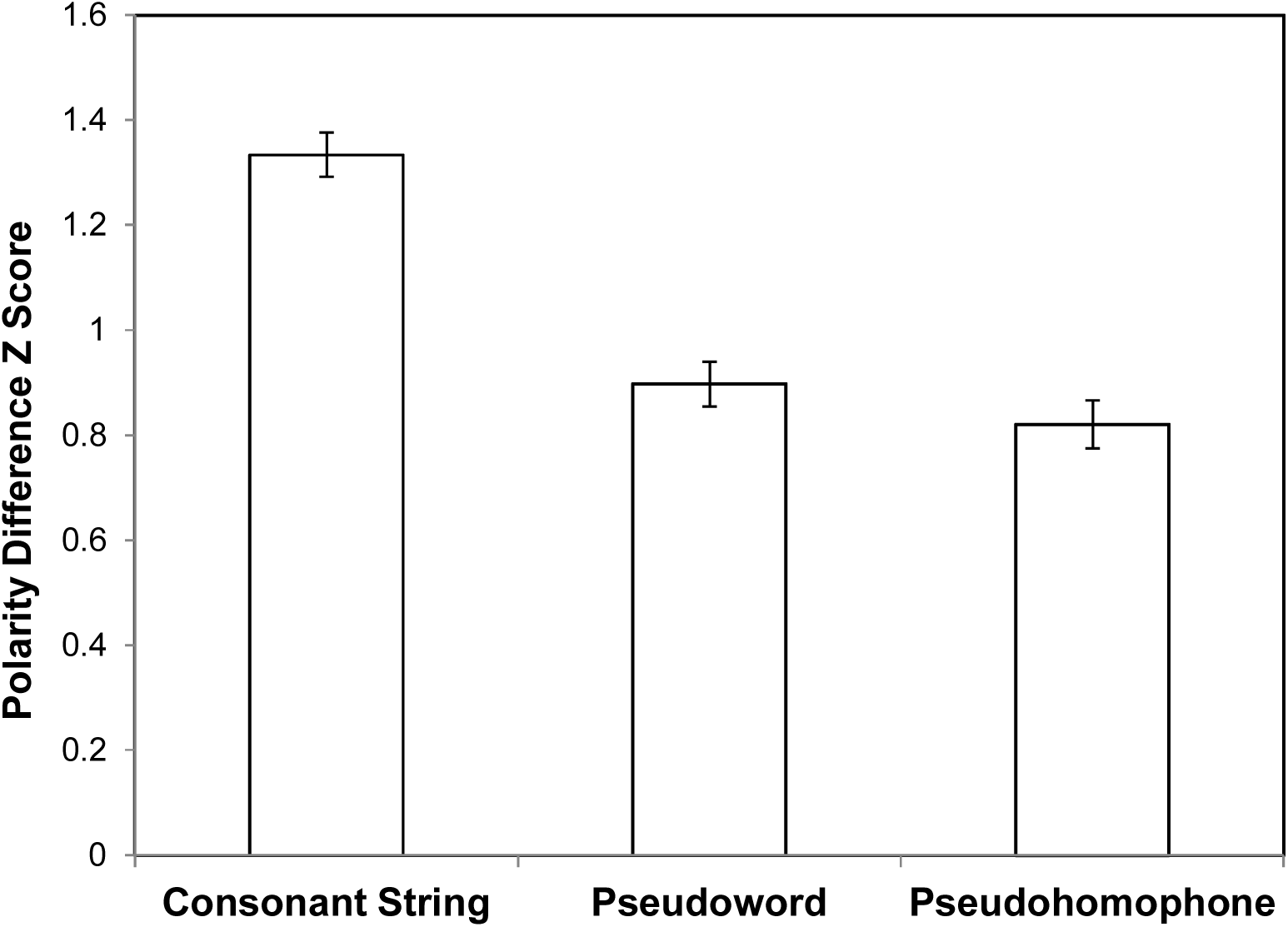

#### 2.3.3. Layer contribution to polarity differences

Figure 6 illustrates the individual contribution from orthographic, phonological and semantic layers to the polarity differences by showing the percentage that each layer contributes to the average differences in the three foil conditions. In the consonant string condition, orthography made the most contribution (48%), while in the pseudoword and pseudohomophone condition, semantics made the biggest contribution of (38% and 36% respectively). T tests revealed that the contribution made by the orthographic layer in the consonant string condition was significantly larger than that in both the pseudoword and pseudohomophone conditions (both *ps* < .05). By contrast, semantic processing was more important in both pseudoword and pseudohomophone conditions relative to the consonant string condition (both *ps* < .05).

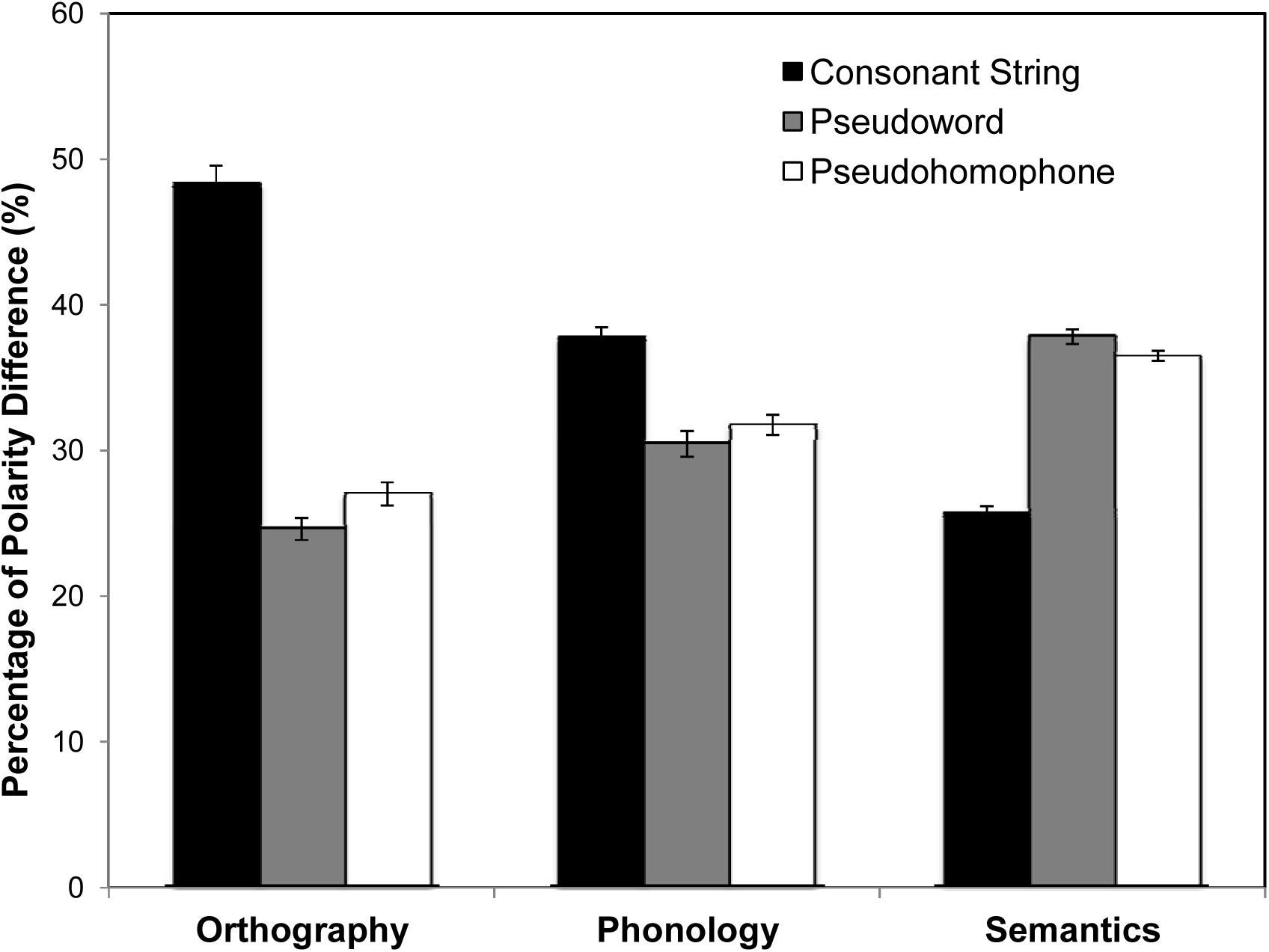

#### 2.3.1. Lexical Decision Criteria

The simulations so far have shown that differences in polarity scores at different layers in the network have the potential to form the basis for lexical decisions. But a crucial question is what criteria should be applied to this measure to allow lexical decisions to be made accurately and quickly? A straightforward approach might be to use a fixed threshold. If the item polarity becomes higher than the threshold, it is a word; otherwise, it is a nonword. However, this is not ideal, because it does not allow for the fact that the polarity measures are time-dependent; early on in processing a relatively low threshold might be appropriate, but later a much higher threshold would be required. Also, this single threshold approach suffers from the problem that nonword decisions can only be made after all processing ticks (when it is known that the threshold will never be exceeded). In reality, nonword decisions can be made very quickly if the nonwords are not very word-like. To avoid these problems we adopted three separate dynamic criteria for word and nonword decisions: (1) word boundary: If at any tick, the polarity score exceeded the average polarity score for nonwords by more than two standard deviations, then a word decision would be recorded; (2) nonword boundary: If at any tick, the polarity score was more than two standard deviations under the average polarity score for words, then a nonword decision would be recorded; (3) minimum activation: the decision can only be made after the polarity reached an active level of 0.8. The last criterion was to ensure that the model could make a decision based on reliable information. The polarity for an item was computed by combining the measures of average polarity for that item in the orthographic, phonological, and semantic layers so that the layers made equal contributions. If at any tick, either of the first two above criteria was met and the polarity score was greater than 0.8, then the lexical decision was assumed to have been made, and the response time was taken as the tick on which the decision was reached. When neither of the criteria was met by the time the last tick was reached, the responses were made based on whether their polarities at the last time tick were closest to the average word and nonword polarities. The response latencies for those items were assigned according to their distance to the average polarity. For the top 10% items, the response latencies were 31 ticks, and for the next 10% items, the latencies were 32 ticks, and so on. The slowest responses for the least 10% items were 40 ticks.

#### 2.3.2. Inverse efficiency

It is worth noting that the cut-off lines for the word and nonwords are arbitrary and could be varied to produce a different speed-accuracy trade-off. To control for these potential differences in a speed-accuracy trade-off, we adopted inverse efficiency as our performance measure. Inverse efficiency is reaction time divided by accuracy, and it is relatively robust to different levels of speed-accuracy trade-off (Roberts, Lambon Ralph, & Woollams, 2010; Roder, Kusmierek, Spence, & Schicke, 2007). To illustrate the effectiveness of inverse efficiency, we tested the model on all the 2,971 words in the training set against a set of nonwords consisting of the same number of monosyllabic pseudowords taken from the ARC nonword database (Rastle, Harrington, & Coltheart, 2002). The length of nonwords ranged from three to seven letters. Different cut-off lines were tested including one standard deviation, two standard deviations and three standard deviations. When the cut-off line of one standard division was used, only 0.2% of word responses did not meet the decision criteria but this increased to 27.2% for the cut-off line of two standard deviations and 69.1% for the cut-off line of three standard deviations. The response latencies of those items were assigned according to their distance to average polarity. Figure 7 shows the distributions of accuracy, response time and inverse efficiency for three different cut-off lines produced by the model. The use of different cut-off lines greatly influenced the distribution patterns of response time while the distribution patterns of inverse efficiency remained similar, indicating inverse efficiency was relatively robust to the selection of different cut-off lines. In the following tests, we opted to use the cut-off line of two standard deviations.

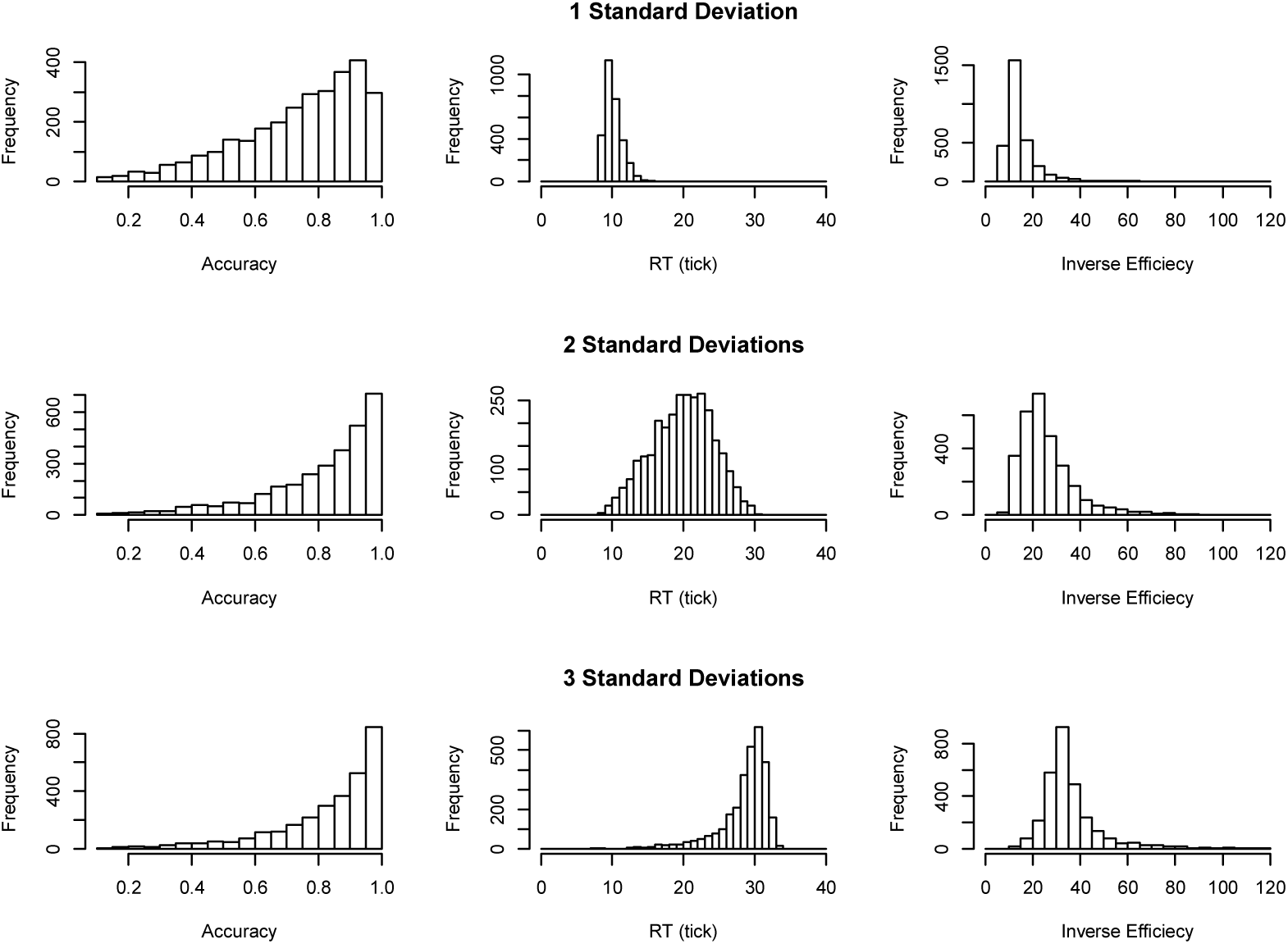

### 2.4. Semantic influences on lexical decision

The key test for the model was to see if the model could produce a graded imageability effect in lexical decision depending on condition difficulties as observed in Evans et al. (2012) where the imageability effect was larger when words were tested in the context of pseudohomophones than pseudowords, and it disappeared altogether in the context of consonant strings as in Figure 8 (panel (a)). We tested the model on the stimuli from Evans et al. Words that were not in the training corpus and their matched nonword items were removed. The remaining 70 words consisted of 35 high- and 35 low-imageability words along with their matched nonword items. Incorrect items and those with inverse efficiencies farther than three standard deviations from the mean were discarded prior to the analyses. A 2×3 repeated measures ANOVA was used to confirm that the model exhibited a similar pattern of results found in the behavioural data. There were reliable main effects of foil condition, F(2, 38) = 54.09, *p* < .001, η_p_ = 0.74, and imageability, F(1, 19) = 6.16, *p* = .023, η_p_ = 0.245, together with the critical significant interaction between imageability and foil condition, F(2, 38) = 8.21, *p* = .001, η_p_ = 0.302. The simple effect analyses showed that the imageability effect was not significant with consonant strings (*p* > .05). There were significant imageability effects in the context of pseudowords, F(1, 19) = 7.43, *p =* .013, as well as pseudohomophones, F(1, 19) = 7.53, *p* = .013, with the effect size slightly larger for pseudohomophones (η_p_ = 0.284) than for pseudowords (η_p_ = 0.281). The results are shown in Figure 8 along with the comparison data from Evans et al. Overall the simulation results show a similar pattern to the behavioural results from Evans et al.

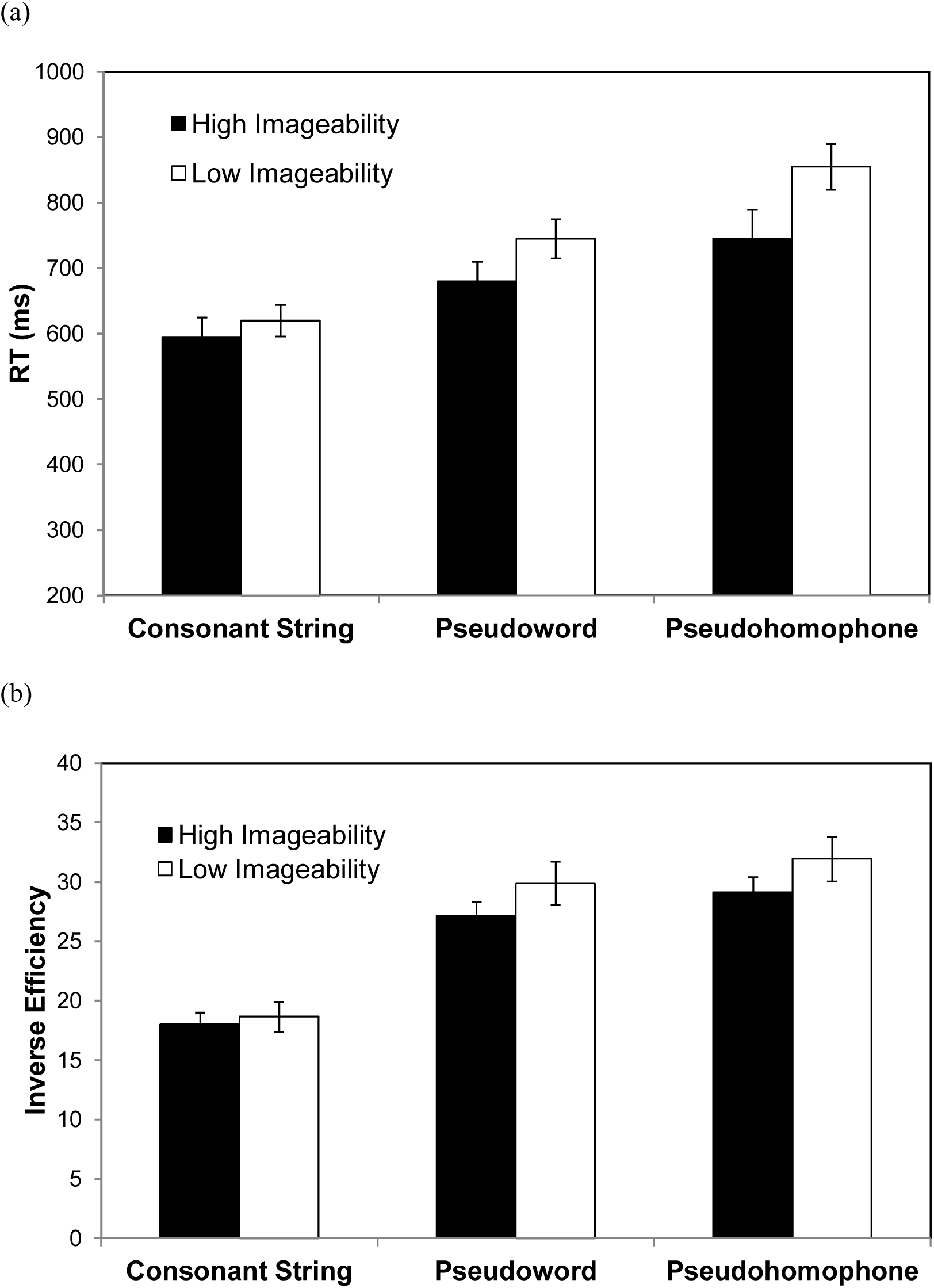

It is clear that the model was able to capture the imageability effect in lexical decision. Nevertheless, all of the semantic vectors for high imageability and low imageability words in the model had the same number of active features. This is very different from previous connectionist simulations (Harm & Seidenberg, 2004; Plaut & Shallice, 1993) that explicitly implement semantic representations with more features for high than for low imageability words. Thus, a question raised is how the imageability effect emerged in the present model. Several studies have reported that words with a high number of shared semantic features generated faster responses than those with fewer shared features when the total number of features are held constant during a lexical decision task and a semantic task (Grondin, Lupker & McRae, 2009), suggesting not all features matter equally. Semantic richness has been proposed to be multifaceted, and each construct could make distinctive influences on lexical processing (Yap, Tan, Pexman, & Hargreaves, 2011; Yap, Pexman, Welsby, Hargreaves, & Huff, 2012). Thus, imageability was likely operationalised in the model in terms of the shared semantic features. If this is the case, we would expect that high imageability words have more shared features than low imageability words. We computed the representational distance matrices for both high imageability words and low imageability words, as in Figure 9. A pair-wise correlation analysis showed that the average cosine distance score for high imageability words (M = 0.867) was significantly lower than that for low imageability words (M = 0.903), t(1188) = 5.45, *p* < .001, confirming that high imageability words contain more semantic features that are shared with others than low imageability words.

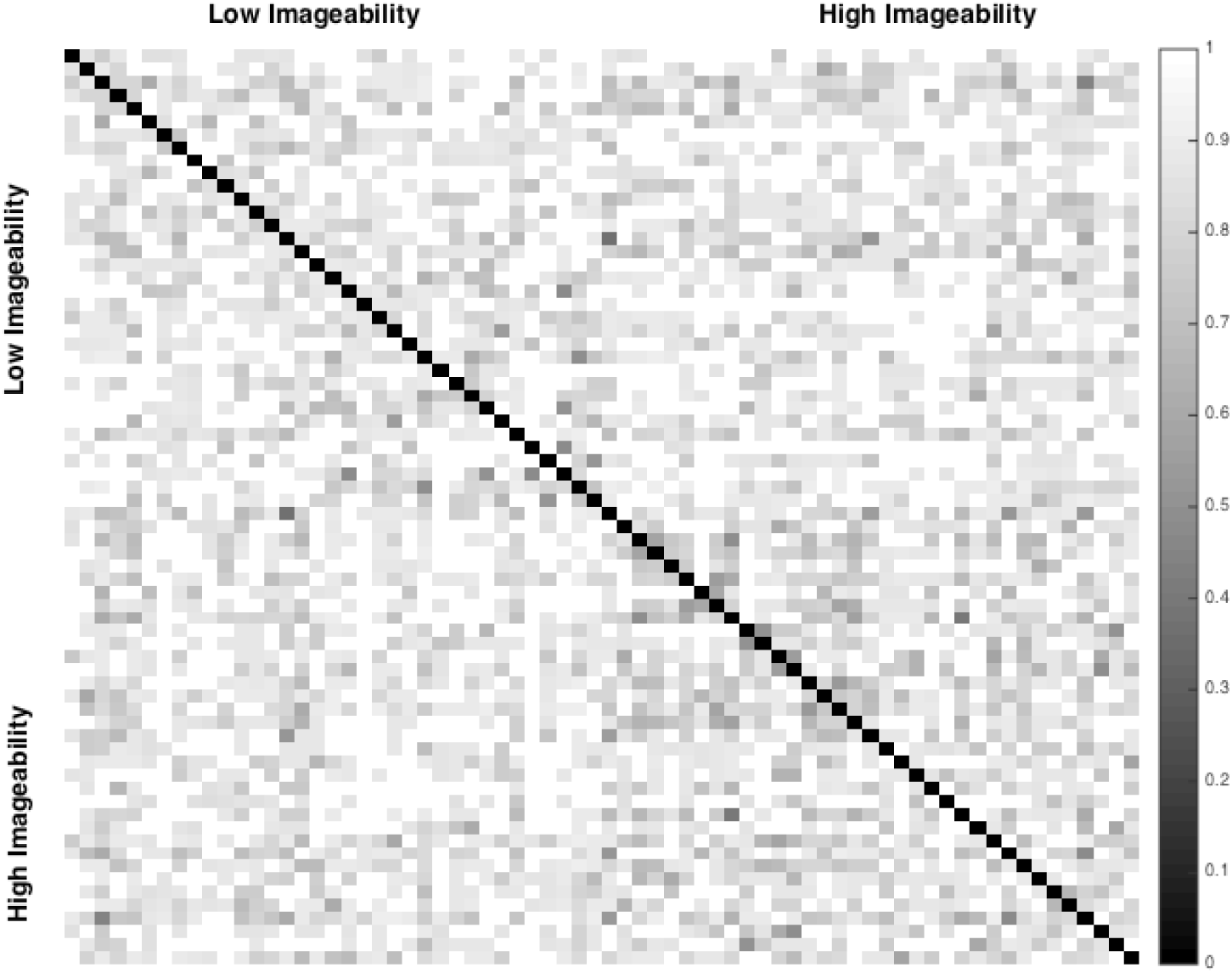

### 2.5. Frequency by consistency effects on lexical decision

The effects of frequency and consistency have been well reported in word naming (Baron & Strawson, 1976; Jared, McRae, & Seidenberg, 1990; Plaut, et al., 1996; Seidenberg & McClelland, 1989; Stanovich & Bauer, 1978; Taraban & McClelland, 1987). However, in lexical decision a small or null effect of consistency is generally reported (Andrews, 1982; Seidenberg, et al., 1984; Waters & Seidenberg, 1985). When studies include orthographically atypical words (i.e., words with low bigram frequencies) (Parkin & Underwood, 1983; Waters & Seidenberg, 1985) or mega studies include a wide range of words (Balota et al., 2004), the effect is likely to be observed. Additionally, the consistency effects in lexical decision also have been reported in patients with semantic dementia (Patterson et al., 2006; Rogers, Lambon Ralph, Hodges, et al., 2004). These results suggest that the effects of spelling-sound inconsistency may not be easily observable when the lexical decision can be made on the basis of perceptual information available prior to the access to phonology, in contrast to when words need to be slowly recognised, or the task requires phonological codes (Waters & Seidenberg, 1985). Here, we tested the model on four sets of stimuli taken from Andrews (1982): high-frequency consistent words, low-frequency consistent words, high-frequency inconsistent words, and low-frequency inconsistent words. Each word set consisted of 20 words except that one high-frequency inconsistent word (*live*) was discarded because it was not in the training corpus. As there were no nonword foils included in supplementary material in the original study, we used the pseudowords taken from Evans et al. (2012).

A 2×2 ANOVA analysis revealed that the main effect of frequency, F(1, 19) = 128.8, *p* < .001, η_p_ = 0.871, was significant. Neither consistency nor the interaction reached a significant level (*ps* > .05). The simulation results are consistent with behavioural findings, suggesting that the consistency effect is not generally observable in factorial designs with a small sample of words. As we shall see in the next section, the effect could be captured by using a regression technique on a wider range of words.

### 2.6. Linear mixed-effect model analyses on lexical decision

We have examined imageability effects and frequency, consistency and their interaction by using factorial analyses. In order to rule out that the possibility of the observed effects being a consequence of specific samples tested, it would be useful to assess if the model could produce similar effects on a larger set of words. Thus, we conducted linear mixed-effect model (LMM) analyses on all the words in the training corpus, akin to the large scale of regression analyses on lexical decision conducted by Balota et al. (2004). It was anticipated that the model should be able to reproduce a range of reading effects in lexical decision including frequency, consistency, neighbourhood size, word length, and imageability as well as the interactions of frequency by word length and of frequency by neighborhood size (Andrews, 1992; Balota et al., 2004; Cortese & Khanna, 2007).

All of the 2,971 words in the training set were tested against the set of nonwords consisting of the same number of monosyllabic pseudowords from the ARC nonword database (Rastle, Harrington, & Coltheart, 2002). The inverse efficiency scores were used as a dependent variable. Item number (one to 2,971) and simulation (one to 20) were included as random effects. A set of psycholinguistic variables was included as fixed effects: frequency (Freq), word length (WL), orthographic neighbourhood size (OrthN), consistency (Con) and imageability (Img). The frequency measure in the model was the frequency of the model’s exposure to each word. Spelling-to-sound consistency score was determined by counting the proportion of words with the same rime that were pronounced in the same way as the target word, and the score was weighted by word frequency (Jared, 1997). Orthographic neighbourhood size was the number of words that could be created by changing one letter in a target word (Coltheart, 1977). Imageability scores for each word were taken from the norm by Cortese et al. (2004). All error responses and outliers (inverse efficiency greater than or smaller than three standard deviations) were excluded. Only words that have known values for all the predictors were used, leaving 48,804 observations for further analyses. The dependent variable was log-transformed because the distribution was skewed, and all the variables were scaled prior to LMM analyses. The correlations between the predictors and inverse efficiency can be found in Table 2.

**Table 2.**
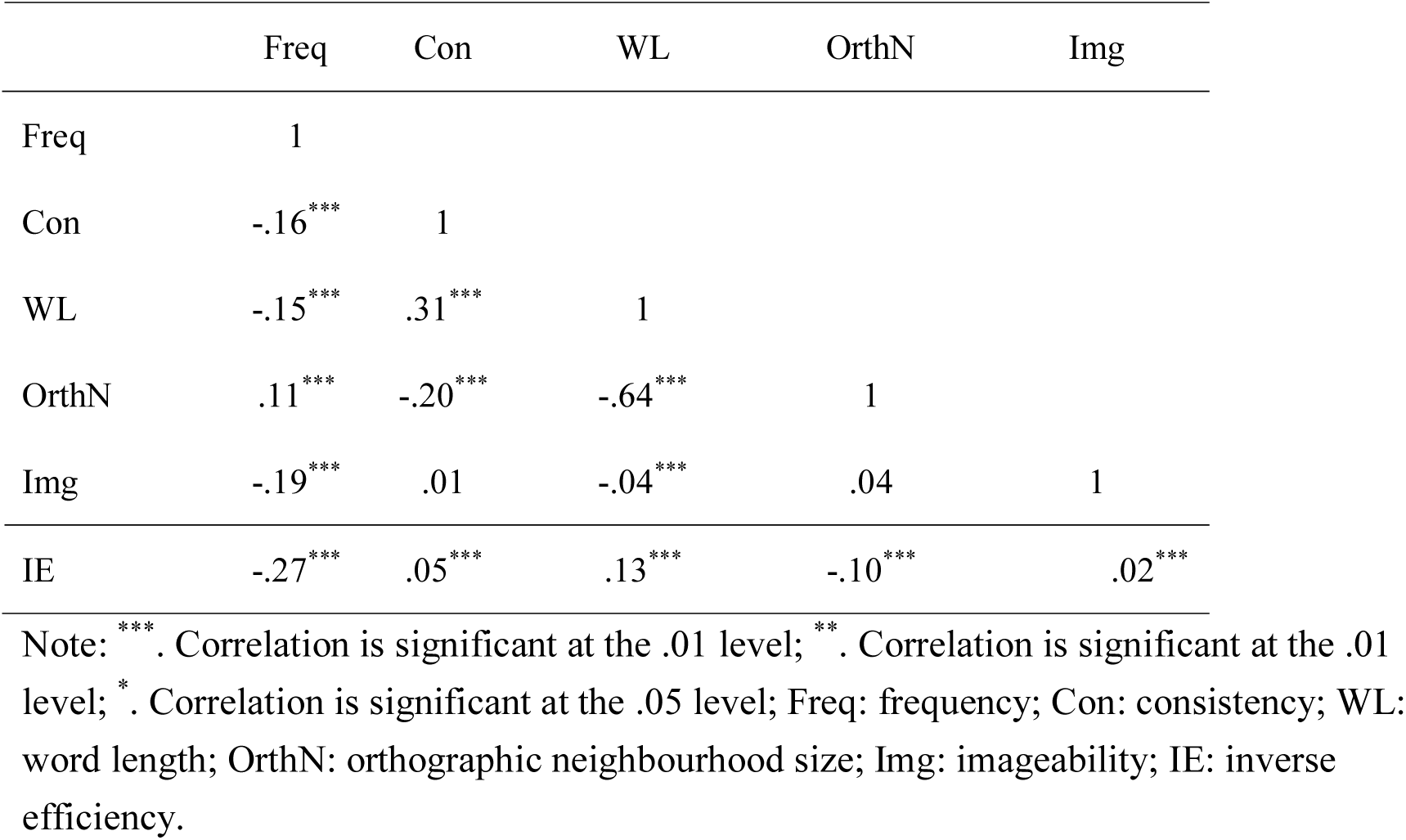
Inter-correlations between all the predictors and inverse efficiency

The LMM result showed that Freq, Con, WL, OrthN and Img all made significant contributions in predicting inverse efficiency (Table 3). The results demonstrated that words that were high in frequency, more imageable and with consistent spelling-to-sound mappings were processed more easily. That was also true for words that were short and had many neighbours.

**Table 3.** Standardised coefficients of the regression model in lexical decision

|  | Estimated | SE | t value |
| --- | --- | --- | --- |
| Freq | -0.351 | 0.011 | -32.53 |
| Con | -0.032 | 0.011 | -3.05 |
| WL | 0.106 | 0.013 | 7.89 |
| OrthN | -0.032 | 0.013 | -2.46 |
| Img | -0.029 | 0.010 | -2.81 |

Note: Freq: frequency; Con: consistency; WL: word length; OrthN: orthographic neighbourhood size; Img: imageability.

To explore the relationship between frequency and word length, all the predictors along with the interaction term were entered into the LMM model. The interaction between frequency and word length was significant, *Estimated* = −.056, *SE* = 0.010, *t* = −5.74. As in Figure 10 (a), the interaction pattern is very similar to that reported in Balota et al. (2004, Figure 12 for young adult), where behavioural RTs for high frequency words are negatively correlated with word length while the RTs for low frequency words are positively correlated with word length. A similar LMM analysis was conducted to examine the relationship between frequency and orthographic neighbourhood size. All the predictors and the interaction between frequency and orthographic neighbourhood size were entered. The result showed that the interaction term between frequency and orthographic neighbourhood size was not significant (*t* = 0.51), though this was found to be significant, *Estimated* = .025, *SE* = 0.012, *t* = 2.06, when the three-way interaction between frequency by neighbourhood size by word length was included, *Estimated* = .028, *SE* = 0.010, *t* = 2.74. In the model, the interaction between frequency and neighbourhood size was modulated by word length. The interaction pattern between frequency and orthographic neighbourhood size as shown in Figure 10(b) is also similar to that reported in Balota et al. (2004, Figure 13 for young adult) and compatible with the finding by Andrews (1989) that the effects of orthographic neighbours sizes were facilitatory for low frequency words while the null or the inhibitory effects were observed for high frequency words (Balota et al. 2004).

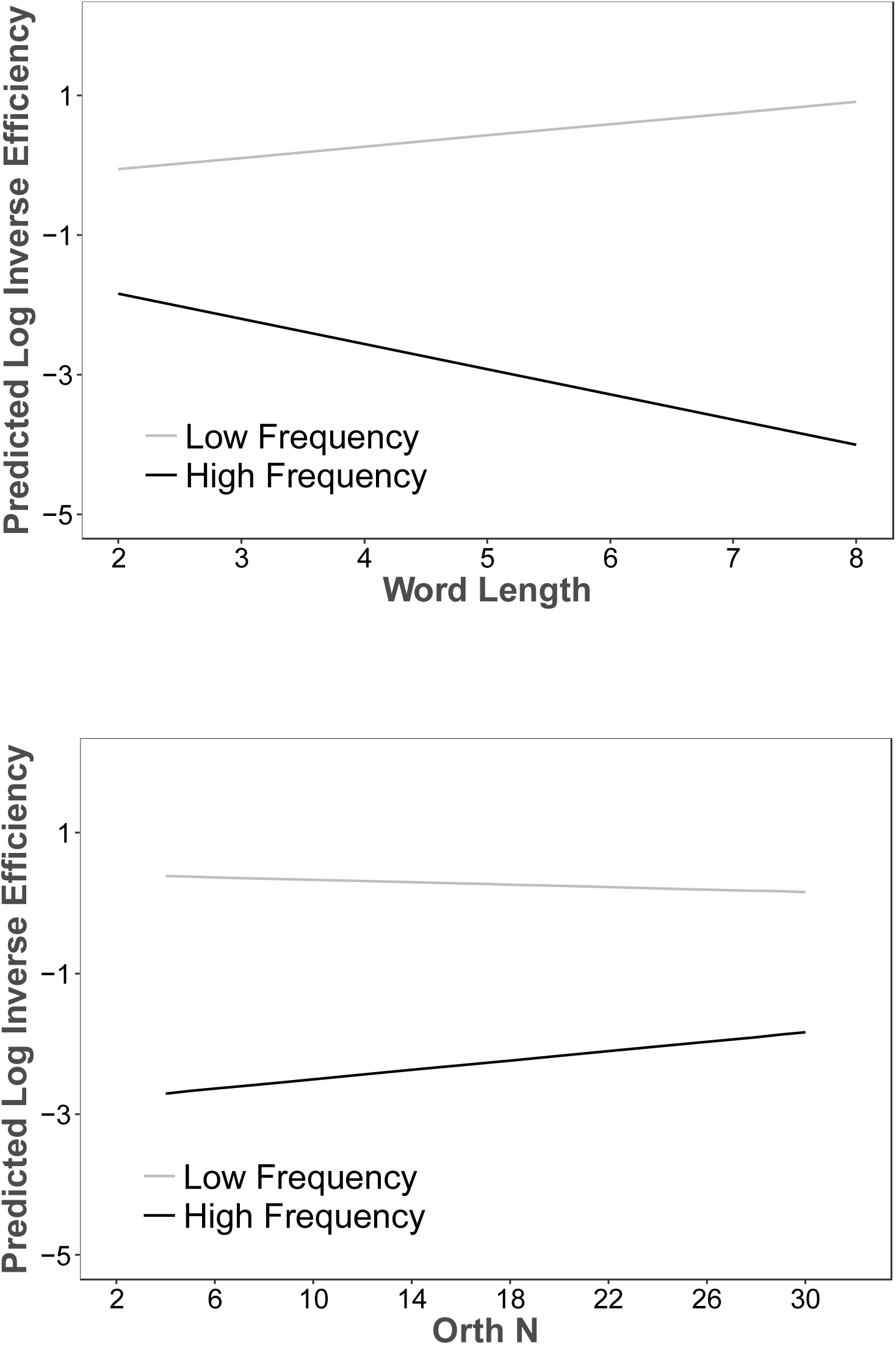

In summary, the present results of the main effects on the model’s lexical decision performance are consistent with those reported by previous behavioural studies (Balota et al. 2004; Cortese & Khanna, 2007). For the interaction patterns, we found that word length was dependent on word frequency (Balota et al. 2004) so was orthographic neighbourhood size (Andrews, 1989, 1992; Balota et al. 2004).

### 2.7. Predicting item-level variance in lexical decision

The issue of predicting item variance of human latencies is one of the most challenging tests for any computational model of reading (Spieler & Balota, 1997). When several influential models of reading aloud including DRC (Coltheart et al., 2001), CDP (Zorzi, Houghton, & Brian, 1998), triangle (Plaut et al., 1996), and CDP+ were tested against on 2,870 words in Spieler and Balota (1997), only 3% to 7% of variance of word naming latencies could be accounted for (Coltheart et al., 2001) This is considerably lower than the variance of human latencies accounted for by the three most important lexical factors (frequency, orthographic neighbourhood size and word length), which is about 21.7% of the variance. The modelling data has been shown to close to the behavioural performance until a new version of the CDP model (i.e., CDP+ model, Perry et al., 2007) has been developed.

While previous studies have focused on the ability of computational models to predict item variance in word naming latencies, to our knowledge, very few studies (e.g., Kello, 2006) reporting item variance in computational models of lexical decision on a large set of words. To address this lack, we tested the present model on its ability to account for the item variance of human lexical decision latencies based on data from the English Lexicon Project (ELP) (Balota et al., 2007). Again, we tested all the 2,971 words in the training set against the same number of pseudowords. We first computed the average lexical decision latencies across simulation runs for each word. We then looked for its lexical decision latencies (z-scored RTs) in the ELP (Balota et al. 2007) where 2,864 words had latencies recorded in the database.

Prior to analysis, outliers (greater than two standard deviations) of the behavioural data were removed, resulting in a removal of 4.2% of words. Inverse efficiencies produced by the model were first log-transformed and then entered into the regression model as a predictor with the behavioural z-scored RTs as a dependent variable. The result showed that the inverse efficiency scores accounted for a significant portion of the z-scored RTs, *R*^2^ = 7.3%, *p* < .001. For comparison, we conducted an additional regression analysis where the three lexical factors including word frequency of Hyperspace Analogue to Language (HAL) (Lund & Burgess, 1996), orthographic neighbourhood size and word length were used as predictors. The result showed that these variables together could predict 42.64% of the variance of z-scored RTs (*p* < .001). The variance predicted by the present model was substantially lower than that predicted by the three lexical factors.

One possibility for this gap in variance is that the frequency range used for training the model is substantially compressed by using a logarithm transformation to reduce the training times. However, this may not be ideal for capturing participants’ item-level frequency effects. To explore this possibility, we conducted an additional regression analysis with log frequency used in the model along with orthographic neighbourhood size and word length as predictors, and with the z-scored RTs as the dependent variable. The result showed that 15.76% of variance of the z-scored RTs (*p* < .001) was predicted, which is higher than the model performance but is much lower than that predicted by the regression model with HAL frequency (42.64%). This suggests that the frequency range used to train the model is at least a potential cause of the discrepancy.

Another issue concerns whether semantic processing contributes much to the item variance accounted for by the model. While the semantic system is crucial for the model to simulate the graded semantic effect in lexical decision, it remains unknown how much added value is contributed by the semantic layer in terms of accounting for item-level variance in human lexical decision data. To test the contribution of semantic processing, we removed all the polarity scores generated from the semantic layer in the model, akin to more focal lesions to semantic processing. We then re-conducted the regression analysis. The result showed that the inverse efficiency scores accounted for a significant portion of the z-scored RTs, *R*^2^ = 4.1%, *p* < .001. It was 3.2% lower than that produced by the full regression model. For comparison, we also conducted two additional regression analyses to test the contributions of phonological and orthographic processing. The results showed that the variance accounted for by the regression model without phonology or orthography was 0.3% or 1.4% lower than the full regression model, respectively. The findings demonstrate that semantic processing indeed greatly helps the model to account for item-level variance in human lexical decision data with orthographic processing the second and phonological processing the least.

## 3. Simulation 2

Having demonstrated that the model can simulate lexical decision by integrating information across visual-orthographic, phonological and semantic processing layers, the next important question is whether damage to those layers would result in general patterns of impaired lexical decision similar to those observed in patients with functionally corresponding reading deficits - namely pure alexia, phonological dyslexia and semantic dementia.

### 3.1. Lexical Decision in Patients with Reading Deficits

#### 3.1.1. Pure Alexia

Pure alexia (PA) is a neuropsychological deficit generally caused by lesions in the left ventral occipitotemporal region (Damasio & Damasio, 1983). The hallmark feature of PA is abnormally strong length effects in word reading times, and it is thought by many to result from damage to visual processing (Arguin, Fiset, & Bub, 2002; Behrmann, Plaut, & Nelson, 1998; Fiset, Arguin, & McCabe, 2006; Roberts, et al., 2010). Despite the visual impairment in PA patients, some are still sensitive to lexical variables such as word frequency (Behrmann, et al., 1998; Johnson & Rayner, 2007; Montant & Behrmann, 2001), regularity (Friedman & Hadley, 1992), orthographic neighbourhood size (Arguin, et al., 2002; Fiset, et al., 2006; Montant & Behrmann, 2001), age of acquisition (Cushman & Johnson, 2011) and word imageability (Behrmann, et al., 1998) in word naming. In terms of lexical decision, some PA patients are able to perform the task above chance (Coslett & Saffran, 1989; Friedman & Hadley, 1992; Roberts, et al., 2010). Their lexical decision performance is modulated by the severity of the condition, word frequency, imageability and nonword type. According to the partial activation account (Behrmann, et al., 1998), some lexical-semantic processing could be activated by bottom-up visual stimuli, albeit to a lesser level relative to unimpaired reading (see the right-hemisphere account by Coslett and Saffran (1994) for a different interpretation of implicit recognition).

#### 3.1.2. Phonological Dyslexia

Patients with phonological dyslexia (PD) are characterized by a relative impairment of nonword reading in the context of better word reading accuracy (Beauvois & Derouesne, 1979; Patterson, 1982). Many studies suggest that the functional locus of phonological dyslexia is disturbance to phonological processing because patients’ reading performance is strongly correlated with their non-reading phonological deficits, and they exhibit the same qualitative performance characteristics on reading and nonreading tasks, including substantial lexicality and imageability effects (Crisp & Lambon Ralph, 2006; Patterson & Marcel, 1992; Rapcsak et al., 2009), although a few isolated studies have reported patients with no significant impairment to phonological processing (Caccappolo-van Vliet, Miozzo, & Stern, 2004; Tree & Kay, 2006). Additionally, consistency effects in word naming are not traditionally associated with phonological dyslexia because it is not often tested. However, when Welbourne & Lambon Ralph, (2007) reanalysed two case series studies of phonological dyslexia data (Berndt et al., 1996; Crisp & Lambon Ralph, 2006), they found that both groups of patients showed a small but reliable consistency effect.

Lexical decision is generally not a critical diagnostic test for phonological dyslexia so, very few studies report it, and when they do, it is as a background test. The findings show that patients have near-perfect performance in judging whether letter strings are words or not (Cuetos, ValleArroyo, & Suarez, 1996; Denes, Cipolotti, & Semenza, 1987). Only when patients are tested with low-frequency words does their performance decline to below typical performance (Dujardin et al., 2011).

#### 3.1.3. Semantic Dementia

Semantic dementia (SD) is characterized by a progressive loss of conceptual knowledge. Several studies have shown that SD patients’ word naming performance is affected by some psycholinguistic properties of words such as frequency, regularity, consistency and imageability (Jefferies et al. 2004; Patterson et al. 2006; Woollams, 2015). Many SD patients also have surface dyslexic reading patterns (Woollams et al., 2007), showing a selective deficit in naming words with inconsistent spelling-to-sound mappings whereas the nonword naming ability is relatively preserved, although there are isolated studies reporting SD patients without significant impairment in reading aloud (e.g., Blazely et al. 2005). In terms of lexical decision, a number of studies have shown that the performance of SD patients is significantly poorer than controls (Benedet, Patterson, Gomez-Pastor, & de la Rocha, 2006; Diesfeldt, 1992; Patterson, et al., 2006; Rogers, Lambon Ralph, Hodges, et al., 2004; although see Coltheart, 2004; Blazely et al., 2005). Importantly, the performance of SD patients depends on the nature of stimuli. Diesfeldt (1992) reported that in a visual lexical decision task, a patient, BHJ, performed well when words were tested against consonant strings but had significant difficulty in distinguishing words from more wordlike nonwords such as pseudowords and pseudohomophones. In the two-alternative forced choice paradigm, the patients were able to judge orthographically typical words from the relatively atypical nonwords, but their performance was significantly impaired in the reverse condition (Rogers, Lambon Ralph, Hodges et al. 2004).

Additionally, several studies have shown an enhanced imageability effect in SD patients (Jefferies et al. 2009; Hoffman & Lambon Ralph, 2011; Hoffman, Jones, & Lambon Ralph, 2013), while others have reported a reversal of the imageability effect (Bonner et al. 2009; Breedin et al., 1994; Yi et al., 2007). The discrepant findings could be due to the selection of stimuli and/or individual differences (Hoffman et al., 2013). When appropriate stimuli are used, most SD patients can process high imageability words more efficiently than low imageability words, because they have more robust semantic presentations. However, most of the enhanced imageability effects are observed when SD patients perform semantic tasks (e.g., synonym judgment and picture-word association). Such tasks require explicit access to semantic knowledge, whereas in lexical decision or word naming the semantic system may be involved, but the task does not require explicit semantic knowledge to be retrieved. A few studies have been conducted to investigate SD patients’ word naming and lexical decision performance, but the findings are inconclusive. For example, Breedin et al. (1994) reported that an SD patient, DM, responded more slowly for concrete than abstract words in an auditory lexical decision task, demonstrating a trend toward reversed imageability effect (*p* < .06). In a case series of SD patients, Reilly, Grossman and McCawley (2006) reported a reversed imageability effect in mild SD patients’ auditory lexical decision performance, but the null effect was observed for more severe SD patients. Moreover, Pulvermüller et al. (2010) found a facilitatory imageability effect in SD patients’ visual lexical decision performance, but the observed effect was potentially confounded with frequency in their study because low imageability words had significantly lower frequency compared to high imageability words. In a recent study, Woollams (2015) demonstrated a reversal effect of imageability in SD patients’ word naming performance, particularly for inconsistent words, suggesting damage to the semantic system might render the related semantic k1nowledge inaccessible or unreliable resulting in an over-reliance on orthographic and phonological knowledge.

In summary, to simulate patients’ data in lexical decision, we first damaged the model by lesioning the functional locus of the corresponding processing layer in the model. For PA the visual and orthographic layers were damaged, for PD the phonological layer was damaged, and for SD the semantic layer was lesioned. The damaged models were retrained for a period of time to mimic the recovery process following brain damage (Welbourne & Lambon Ralph, 2005, 2007; Welbourne et al. 2011). After retraining, we tested the damaged models on the two sets of data used in Simulation 1 so that we can compare the performance of the damaged models with the intact model. The tests included: (1) frequency and consistency effects from Andrews (1982); (2) imageability and foil effects from Evans et al. (2012).

In line with patients’ behavioural data, we predicted that the PA model would show generally impaired lexical decision performance compared to the intact model but with some sensitivity to lexical-semantic properties of words. Importantly, the performance of the PA model would be strongly modulated by the foil conditions because orthographic processing was more demanding for word-like nonwords than for consonant strings. Though the difference in the contexts of pseudowords and pseudohomophones could be small as both fitted the orthographic structures of words.

For the PD model, we predicted slightly impaired performance with frequency, consistency and imageability effects similar to those observed in the intact model, because phonological processing is relatively less crucial for lexical decision. For the SD model, we predicted strong consistency effects. Regarding imageability effects, the evidence in the literature is equivocal, but we expected that in the presence of word-like foils we might be able to detect a reversed imageability effect in line with Woollams (2015).

### 3.2. Method

The method to simulate PA and PD patient types was similar with the only difference being the location of the damage and the amount of retraining required for the model to recover to a stable performance level. For PA damage, we randomly removed 90% of the links connecting to or from the HO layer coupled with 90% of the links into or out of the connected control units. The network was retrained for 300,000 epochs. For PD damage, the network was damaged by randomly removing 90% of the links into and out of the phonology layer together with 90% of the links into or out of the connected control units. This time the network was retrained for 400,000 epochs. To capture some variations within patients, we analysed the average model performance across five consequent time points toward the end of training. Specifically, we analysed the average performance of the PA model from 260,000 to 300,000 epochs and PD model from 360,000 to 400,000 epochs in steps of 10,000 epochs. Semantic dementia is unlike the other two deficits as it is the result of a progressive disorder (Hodges, Patterson, Oxbury, & Funnell, 1992). Following Welbourne and colleagues (2005, 2007, 2011), we simulated it by repeatedly interleaving very mild damage and retraining. We randomly removed 0.8% of the links into or out of the semantic layer together with the links into or out of the connected control units and then trained the network for one epoch. This process was repeated 400 times.

### 3.3. Results

All of the damaged models were tested in the same way. For the frequency by consistency test, a 2 x2 ANOVA (frequency: high and low; consistency: high and low) was conducted on inverse efficiency scores. Note that we also reported accuracy data in Table 4 for additional information. However, as our demonstration of speed and accuracy trade-off patterns when using arbitrary cut-off lines (Figure 7), we did not conduct analyses on accuracy data. Instead, we decided to stick to the analyses on inverse efficiency to avoid the potential issue. For the imageability by foil test, a 2×3 ANOVA (imageability: high versus low; foil type: consonant string, pseudoword or pseudohomophone) was conducted. The performance of the PA, PD and SD models, as well as the intact model on the two lexical decision tasks, are summarised in Tables 4 and 5.

**Table 4.**
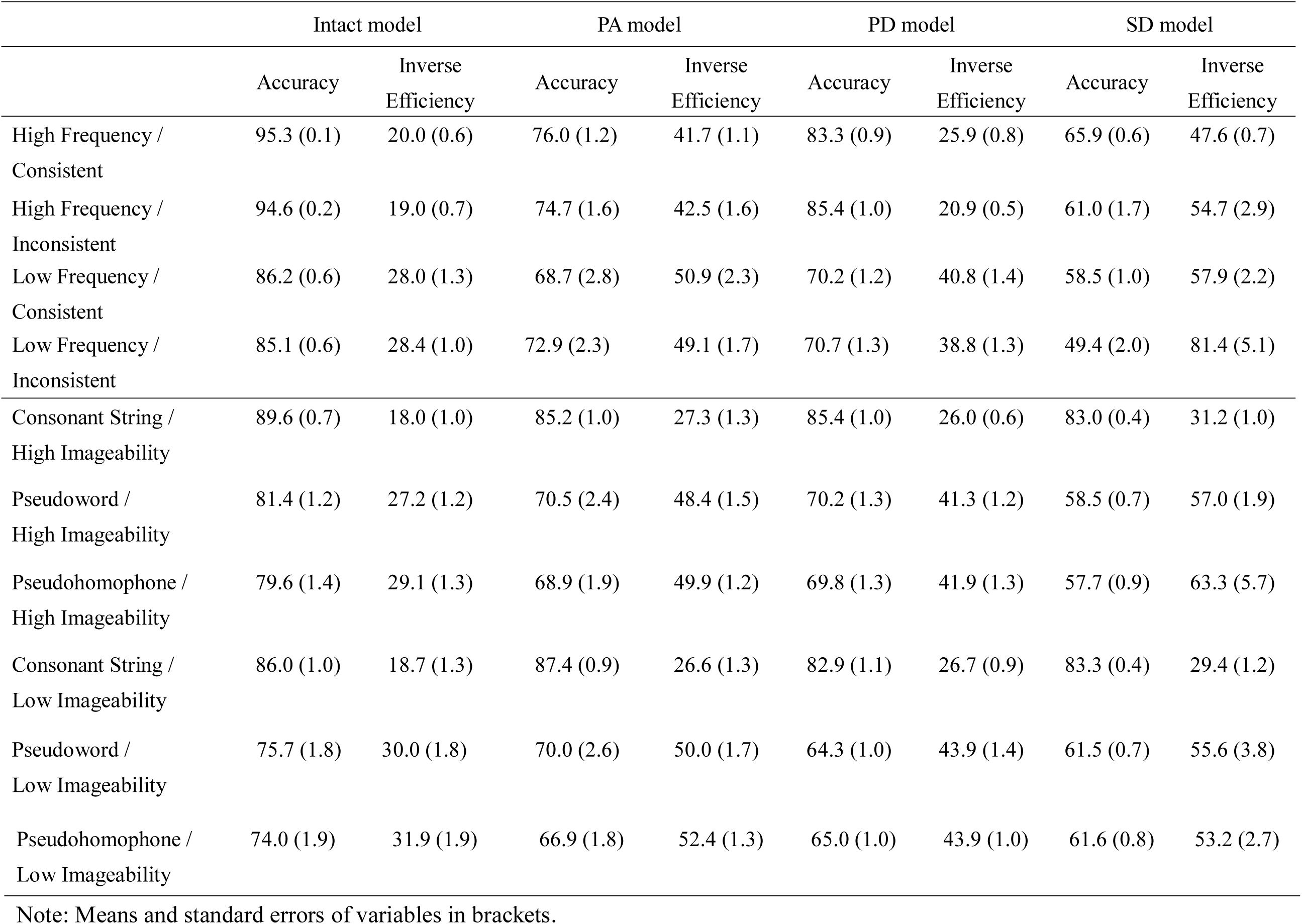
Descriptive results of the intact and the damaged models on the three reading tests in lexical decision.

As indicated in Table 4, the performance of the PA model was generally impaired compared to the intact model. The statistical results in Table 5 showed that the PA model produced significant frequency effects, whereas both the consistency effect and its interaction with frequency were not statistically reliable. For the imageability by foil test, the main effect of imageability was not significant but, more importantly, the main effect of foil type was significant. There was a significant interaction between imageability and foil type. An analysis of simple effects showed that the imageability effect was obtained in the pseudohomophone condition where high imageability words (M = 49.9) were processed more efficiently than low imageability words (M = 52.4), F(1, 19) = 5.25, *p* = .034, η_p_ = 0.22. This result demonstrated semantic processing is involved in judging words against the most word-like foil condition.

**Table 5.**
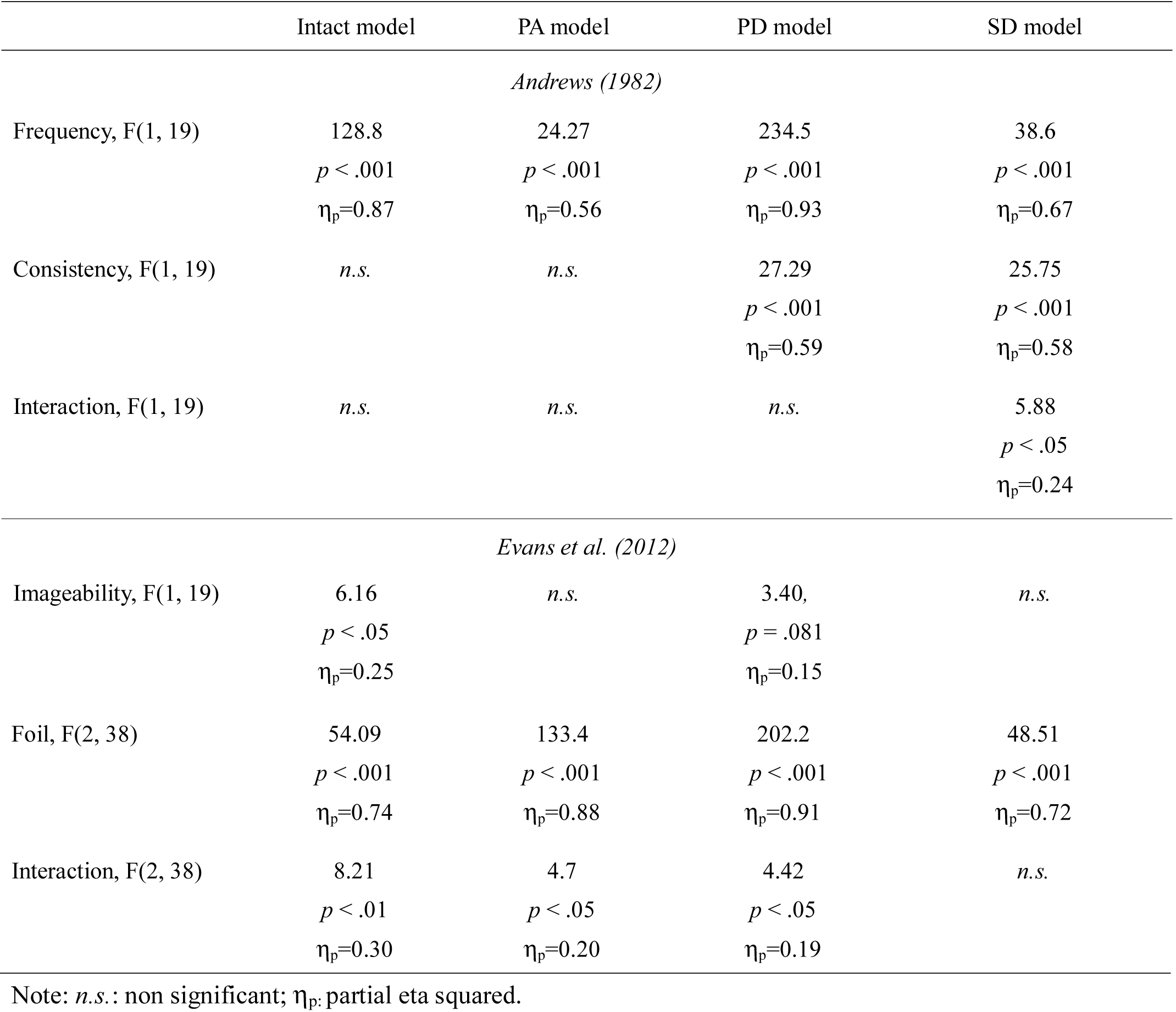
Statistical results of the intact and the damaged models on the two reading tests in lexical decision.

For the PD model, there were significant main effects of frequency and consistency but the interaction was not. The PD model produced higher average inverse efficiency for high consistency words than for low consistency words. Further analyses revealed that the reversal of the consistency effect was caused by semantic processing. Re-running the analyses without semantic contribution resulted in a facilitatory consistency effect, F(1, 19) = 8.4, *p* < .01, η_p_ = 0.31, where high consistency words were processed more efficiently than low consistency words. For the imageability by foil test, the effect of foil was significant, whereas the effect of imageability approached significant (*p* = .081). Importantly, there was a significant interaction. An analysis of simple effects revealed that the imageability effect was not significant in the consonant condition (*p* = .36), while the effect approached to significant in both the pseudoword condition, F(1, 19) = 4.31, *p* = .052, η_p_ = 0.19, and in the pseudohomophone condition, F(1, 19) = 3.71, *p* = .069, η_p_ = 0.16, with high imageability words being processed more efficiently than low imageability words. These results suggest some involvement of semantics in the word-like foil conditions.

With regard to the SD model, the lexical decision performance was generally poor. However, the SD model produced significant effects of frequency, consistency, and their interaction in lexical decision. In particular, the model’s performance was strongly affected by spelling-to-sound mappings because the damaged semantic system could not effectively support recognition of inconsistent words. For the imageability and foil test, although the average inverse efficiency for high imageability words (50.5) was numerically larger than that for low imageability words (46.1), the effect did not research significance, *p* = .16. There was also a significant effect of foil condition whereas its interaction with imageability was not significant.

## 4. General Discussion

The primary aim of this paper was to investigate the processing underlying lexical decision tasks by developing a large-scale recurrent model containing visual, orthographic, phonological, and semantic components. Simulation 1 described how the model was developed by combining oral language and reading training. The model was then used to make lexical decisions by combining information across different processing components, and to explore what contribution each processing component made to the decisions in different contexts. In particular, we were interested in exploring the role of semantics in lexical decision. Based on the measure of polarities at the core processing layers (orthography, phonology and semantics), the model was able to produce the differential semantic effects corresponding to the different nonword foils. The model was also able to capture a range of reading effects including frequency, consistency, word length, orthographic neighbourhood size, and imageability in lexical decision. When information from selected layers was damaged, the model produced similar patterns as observed in patients who have difficulty in accessing that particular type of information. These results demonstrate the ability of the model to account for normal and impaired lexical decision.

### 4.1. Comparison with Other Recurrent Models

In the literature, several large-scale connectionist models of single word reading have addressed the challenge of modelling temporal dynamics in a recurrent framework (Chang, Mohaghan, & Welbourne, 2019; Chang & Monaghan, 2018; Harm & Seidenberg, 1999, 2004; Plaut, et al., 1996; Monaghan et al., 2017). The reading model developed by Harm and Seidenberg (2004) probably provides the most complete simulation of human reading behaviours. Their model demonstrates the division of labour between the phonological and semantic processes, illustrating the cooperative and competitive nature of the reading system. When compared with Harm and Seidenberg (2004)’s model, the present model can be considered as an extension of the same framework. The model here was developed according to the same connectionist principles and was also a large-scale continuous recurrent model with embedded pre-existing phonological and semantic knowledge. However, the present model is different from the previous work in four important aspects:

1. The model includes an additional visual processing component and starts the process from vision to orthography before splitting to both phonology and semantics. The orthographic component can be considered as a layer of hidden units (i.e., OH layer in the model) and the representations are allowed to develop through the acquisition of reading skills as in humans. This implementation is supported by evidence from our previous work (Chang, et al., 2012a), which demonstrated that parallel models can account for the interaction pattern between word length and lexicality observed in reading aloud (Weekes, 1997), provided that orthographic representations are not pre-defined and can be learned over the time course of training. This additional visual processing component also allows the model to simulate the behavioural patterns observed in patients with visual-orthographic deficits.
2. The semantic representations are generated differently and include negative as well as positive components. In Harm and Seidenberg (2004)’s study, the semantic features are generated by using WordNet (Miller, et al., 1990) online database. Each feature is interpretable because it is obtained from the lexical relations among a number of synonymous sets. However, the number of semantic features for a word generated from WordNet will vary with the word set, and the range is rather wide. For example, the number of semantic features in the Harm and Seidenberg (2004)’s study ranges from 1 to 37. To ensure that the semantic activations for a word are mainly affected by its lexical properties rather than the variable-length coding scheme, the semantic representations in the present model are based on the co-occurrence statistics method (Chang et al., 2012b), which provides a fixed-length code that carries critical features for each item. In addition to the conventional use of positive components indicating what attributes an item have, this set of semantic codes includes negative components carrying information about what attributes the item does not have.
3. The architectures of two reading models are slightly different. In Harm and Seidenberg (2004)’s model, there are additional two sets of connections for mappings from orthographic units directly onto phonological units and semantic units to facilitate performance and improve the nonword reading. In the light of our research into the importance of control units for training a model with deep structures (see Appendix A), it may be reasonable to assume that their training issues were largely caused by the pre-trained weights used for the simulation of the pre-existing phonology and semantic knowledge. Adding the direct connections from orthography to phonology and semantics is likely to alleviate this problem as it effectively reduces the network depth. Compared to their model, the present model has deeper structures (i.e., two additional layers for visual processing) and this substantially hinders the learning of the model without control units. Using local control units for each layer with inputs that mirror the inputs of the layer they are controlling is a very effective way of dealing with this problem.
4. The current investigation explored both the nature of ‘normal’ lexical decision and also across three types of acquired dyslexia. Both sources of empirical data have strongly influenced theories and models of reading behaviour. Accordingly, it is interesting and important to test whether implemented models of lexical processing can capture both sources of behavioural data.

### 4.2. Normal Lexical Decision

The most significant contribution from this paper is that the model extends the domain of connectionist models by explicitly tackling a range of lexical decision related phenomena including effects of foil type, frequency, imageability, consistency, word length and orthographic neighbourhood size. There is considerable evidence showing that distinguishing words from consonant strings is generally fast and accurate (Evans, et al., 2012; James, 1975; Ratcliff, et al., 2004; Shulman & Davison, 1977). It is likely that the decision can be made mostly on the basis of visual and/or orthographic information (e.g., Grainger & Jacobs, 1996), only when the decisions become difficult deeper processing would be required (Plaut, 1997; Seidenberg & McClelland, 1989). This is supported by the result shown in Figure 6, where in the context of consonant strings the orthographic processing layer contributed the most to the polarity differences while in the context of more word-like nonwords the semantic layer was critical.

The critical test for the present paper was to see whether the model could account for the differential semantic effects in lexical decision when words were tested with different types of nonwords. The extent of involvement of semantics in lexical decision is controversial (Balota & Chumbley, 1984; Chumbley & Balota, 1984; Coltheart, et al., 2001; Dilkina, et al., 2010; Plaut, 1997). This is mostly because semantic information was initially thought to become available only after lexical access has completed and that lexical access is the only process in the lexical decision task (Becker, 1980; Collins & Loftus, 1975; Forster, 1976; Morton, 1969). This means that there would be no semantic influences on lexical decision. This view has greatly influenced the subsequent theoretical and computational models of lexical decision focusing on orthographic processing (Coltheart, et al., 1977; Coltheart, et al., 2001; Grainger & Jacobs, 1996; Perry, Ziegler, & Zorzi, 2007).

However, evidence from behavioural (e.g., Balota & Chumbley, 1984; Chumbley & Balota, 1984; Balota et al., 2004; Corteness & Khanna, 2007), neuroimaging (e.g., Binder et al., 2003; Hauk et al., 2006; Woollams et al., 2011) and patient studies (e.g., Patterson et al., 2006; Rogers, Lambon Ralph, Hodges, et al., 2004) have suggested that semantic processing is involved in lexical decision albeit to different extents dependent on the properties of word stimuli and the foil types (e.g., Degroot, 1989; James, 1975; Joordens & Becker, 1997; Evans et al., 2012). In particular, Evans et al. (2012) showed that the semantic effect was stronger with pseudohomophones than with pseudowords and than with consonant strings. This pattern was replicated by the model. The simulation results demonstrated that there was a null effect for consonant strings and the effect size of the imageability effect was larger for pseudohomophones than for pseudowords, albeit the difference between pseudohomophones and pseudowords was not as strong as those observed in Evans et al.’ data. Additionally, as shown in Figure 8, the magnitude of semantic effects increased when nonwords became more wordlike. In fact, the graded semantic effects are predicted directly by the time dynamics of polarity difference analyses. Of particular note is that the panels (a) and (c) of Figure 4 shows polarity differences between words and three different types of nonword foils in the orthographic and semantic layers. On average, the polarity differences in the semantic layer were larger than the differences in the orthographic layer; however, the reliable differences in the semantic layer emerged relatively late compared with that in the orthographic layer. These results indicated the importance of semantic access for difficult decision conditions at a later time while for the easier conditions the decisions could be made very quickly based on orthographic information or along with phonological information. Collectively, the findings are consistent with a distributed view of lexical decision, which proposes that both orthography and semantics are important for lexical decision (Dilkina, et al., 2010; Seidenberg & McClallend, 1989; Plaut, 1997). This is in contrast with the localist view arguing for no or little involvement of semantics in lexical decision (Coltheart, et al., 1977; Coltheart, et al., 2001).

In addition to the exploration of the graded semantic effects in lexical decision, linear mixed-effect model analyses on model performance demonstrated that the model was able to account for a range of standard reading effects in lexical decision including frequency, consistency, word length, orthographic neighbourhood size, and imageability. The results also demonstrated that both word length and orthographic neighbours influenced lexical decision performance particularly for low frequency words (Andrews 1989; Balota et al. 2004). These findings are in accordance with the key behavioural effects of psycholinguistic factors influencing lexical decision revealed in behavioural studies using large-scale regression analyses (Balota et al. 2004; Cortese & Khanna, 2007).

### 4.3. Impaired Lexical Decision

Different types of acquired dyslexia including pure alexia (PA), phonological dyslexia (PD), and semantic dementia (SD) were simulated by damaging the correspondent functional impairment in the model and then following by a period of retraining. The results demonstrated that the models were able to reproduce their impaired lexical decision performance.

Pure alexic patients are characterised by a general visual-orthographic impairment. Consequently, their performance on the visual related task would be disrupted. Compared to the intact model, the PA simulations showed substantially impaired lexical decision performance. The performance was strongly affected by the foil conditions in which the PA model could better discriminate words from consonant strings than from word-like nonwords, presumably due to differential demands in orthographic processing. In addition, although explicit word recognition may be difficult, some PA patients have shown sensitivity to frequency, imageability, and nonword type in lexical decision (Coslett & Saffran, 1989; Friedman & Hadley, 1992; Roberts, et al., 2010). According to the partial activation account (Behrmann, et al., 1998), PA patients’ performance may be supported by the partially activated lexical semantic system. This is consistent with the PA simulations, demonstrating a reliable frequency effect. The PA model also showed some degree of imageability effects, particularly when semantic information was critically demanded in the most word-like (i.e., pseudohomophones) condition. Note that the imageability effect in PA patients’ reading is not universally observed (Howard, 1991; Patterson & Kay, 1982; Roberts et al. 2010). One factor affecting PA patients’ recognition performance, which is not the focus here, is related to the severity of the reading deficit. For instance, Roberts et al. (2010) demonstrated that the sensitivity of patients with pure alexia to word imageability was regulated by reading severity. In either very mild or too severe cases, PA patients would not show the imageability effects. This is because for mild PA patients orthographic information could still be reliable for effectively lexical decision while for severe PA patients orthographic activations may not be able to spread to the semantic layer. Collectively, evidence from the current simulation results and the case-series studies seem to suggest the imageability effects in PA patients would be moderated by both foil condition and severity. Future studies can be conducted to vary the levels of visual-orthographic damage to investigate PA patients’ lexical decision in the contexts of different foil types.

As for the PD model, the lexical decision performance was relatively unaffected compared to the other damaged models. The simulation results are congruent with previous studies of phonological dyslexia (Cuetos, et al., 1996; Denes, et al., 1987; Dujardin, et al., 2011), reporting patients’ little difficulty in making lexical decisions, and suggesting that phonology has a minimal role in lexical decision. One divergent result is that the PD model showed inconsistency words were processed more efficiently than consistency words. Within the connectionist framework of reading, inconsistency words are mainly processed via the semantic pathway from orthography to semantics, while consistency words are mainly processed via the phonological pathway from orthography to phonology (Plaut et al., 1996). When the phonological system is extensively damaged, inconsistent words may be less affected and show processing advantage. This explanation is supported by our additional analysis, demonstrating that a facilitatory consistency effect can be obtained with the removal of semantic contribution. However, as PD patients are not generally examined for the consistency effect in lexical decision, whether PD patients with more severe and extensive lesions would show a similar pattern to the simulation result warrants further investigation.

The simulations of semantic dementia in lexical decision showed the strong frequency and consistency effects. In particular, the model had great difficulty in processing low-frequency inconsistent words. The result is compatible with the behavioural data observed in SD patients (Jefferies et al. 2004; Patterson et al. 2006), indicating the importance of semantic processing in inconsistent word reading. Moreover, SD patients perform relatively well on rejecting consonant strings compared to word-like nonwords. (Diesfeldt, 1992). A similar pattern is also observed in the SD model in which the decisions on words in the context of consonant strings was more efficient compared to that of words in the contexts of pseudowords and pseudohomophones as a consequence of preserved orthographic and phonological information but unreliable semantic information. Regarding the imageability effect in the SD model’s lexical decision, the model produced a null effect with numerically lower processing efficiency for high imageability than low imageability words, which is consistent with Reilly et al.’s (2006) findings of moderate to severe SD patients in the auditory lexical decision task. Previous research has shown an enhanced imageability effect for SD patients in the semantic tasks (Jefferies et al., 2009; Hoffman & Lambon Ralph, 2011; Hoffman, Jones, & Lambon Ralph, 2013) whereas a null or reversed effect in the lexical decision and word naming tasks (Breedin et al., 1994; Reilly et al., 2006; Woollams, 2015; but see Pulvermüller et al., 2010). The divergence seems to reflect the task difference concerning whether or not explicit semantic knowledge is required. In lexical decision, decisions can be made on the integrated information of orthography, phonology and semantics. When the information from the semantic system is damaged and becomes unreliable, greater reliance on orthographic and phonological information might be expected. A reversal effect of imageability might be interpreted as being due to greater use of the mappings between orthography to phonology for low imageability words (Woollams, 2015). The severity of semantic dementia could also be a determinant factor (Reilly et al., 2006). However, this would require more systematic investigations on a large-scale case series of SD patients before consensus can be reached.

### 4.4. Comparison with Other Models of Lexical Decision

There are several existing lexical processing models such as the multiple read-out model (MROM) (Grainger & Jacobs, 1996), the dual-route cascade (DRC) model (Coltheart, et al., 2001) and the connectionist dual process (CDP+) model (Perry, et al., 2007) that all share the same lexical processing and decision mechanisms. These models can simulate several effects in lexical decision including the frequency effect, the neighbourhood size effect and the strategic influences on lexical decision by flexibly adjusting decision criteria. Their results are almost all based on orthographic processing with little attention to other processing components, in particular, the semantic system. Thus questions as to how these models could implement a localist view of semantics in lexical decision - presumably either by implementing feedback connections from semantics to their orthographic lexicon (Coltheart, et al., 2001), or adjusting the decision criteria corresponding semantic processing to account for the semantic influences on lexical decision - remain unclear and have not been implemented. Moreover, the MROM model and its variants adopt the interactive activation (IA) model as a visual processing component. Hence, they are inevitably limited to deal with a particular small set of words as they lack a learning mechanism. In contrast, the visual input in the present model is much more flexible and could support learning large-scale word sets.

The present model is compatible with the idea of using accumulated information for lexical decision (Busemeyer & Townsend, 1993; Norris 2006; 2009; Ratcliff et al., 2004; Usher & McClelland, 2001, 2004). In fact, the polarity of the units is a measure of the quality of information produced from the model in response to the stimulus. According to Ratcliff et al. (2004), the decision is made when the accumulation of information over time reaches the appropriate boundaries. The rate of accumulation is assumed to vary as a function of the properties of words and nonwords. However, these types of diffusion model are designed to be decision-making modules only, which means they do not simulate the processes involved in word recognition. We go further to demonstrate the present model can simulate human lexical decisions by using the accumulation of information (i.e., polarity scores) that is integrated across different processing layers over time in the parallel distributed model of reading. This information can directly reflect the different lexical properties of the stimulus. Thus the present model also offers an opportunity to simulate how the effects of lexical properties of word stimulus change with reading experience as reflected in exposure and vocabulary knowledge (Monaghan, Chang, Welbourne, & Brysbaert, 2017; Yap, Balota, Sibley, & Ratcliff, 2012; Yap et al., 2015).

Previous models of word reading have tried to simulate item-level variance in human word naming performance (Coltheart, 2001; Perry, et al., 2007; Plaut et al. 1996) but fewer attempts have been made to capture item-level variance in human lexical decision performance (Kello, 2006). To bridge this gap, we tested the model’s performance in predicting item-level variance in participants’ RTs collected by the ELP (Balota et al. 2004) on over 2,500 English words. The result showed that the model was able to account for 7.3% of the variance of the z-scored RTs in lexical decision, which is slightly higher than that achieved by most previous models in accounting for item-level variance in human word naming performance on a similar test set except the CDP+ model. Further analysis demonstrated that the frequency effect in item-level lexical decision latencies may not be fully captured by the model. This could be improved by training the model to develop a larger vocabulary, and using natural rather than log frequency to determine the probability of presenting each word. While these changes would allow the model to be exposed to a broader range of frequency, they would also considerably increase the training time. The present version of the model represents a sensible compromise between these two considerations.

Finally, the measure of polarity used in Plaut (1997) study and the present model, in a sense defined in equation (3), can be thought of as representing the task-related activity of units in the model. This is potentially interesting because it might also be used as a proxy of the Blood Oxygen Level Dependent (BOLD) signal in neuroimaging studies. Higher polarities incur a higher processing cost, which might map onto the data from fMRI imaging studies. Most existing reading models have been developed to simulate behavioural data (Coltheart, et al., 2001; Harm & Seidenberg, 2004; Plaut, et al., 1996; Seidenberg & McClelland, 1989) and some have been used to account for electrophysiological data (Cheyette & Plaut, 2017; Laszlo & Plaut, 2012; Rabovsky & McRae, 2014); however, relatively little research has been done to account for data from neuroimaging studies of printed word processing. The analyses of polarity differences at each layer in the model could potentially reveal the relative contribution of these layers to the lexical decision tasks. These differences might be expected to relate to the nature of differential processing in the brain regions that support reading. For example, the model was more reliant on semantic information when the nonword foils were pseudohomophones rather than consonant strings. These results resemble the differential brain activation seen in left anterior temporal lobe, which has been associated with semantic processing (Rogers, Lambon Ralph, Hodges, et al., 2004) for lexical decision tasks (Woollams, et al., 2011) and for reading words with atypical pronunciations (Hoffman et al., 2015). Exploring this would require further investigations beyond the scope of the current study.

### 4.5. Limitations and Future Directions

The present model has demonstrated a range of important phenomena in both normal and impaired lexical decision. It should be acknowledged that there are some limitations to this study. First, the model’s decision criteria based on the average word and nonword polarity scores were static rather than dynamic. One potential issue in relation to human readers’ lexical decisions is that the average word polarity scores could be considered as an expectation for words that participants have already built through their experience with words, at least to some extent, prior to the experiment; however, the participants could not have built such as an expectation for nonwords (i.e., the average nonword polarity score) before they actually encounter them. Thus, it might be anticipated that the decision criteria used by the participants in the earlier trials of the experiment may be slightly different from those used in the later trials where the decision criteria may gradually become steady. However, in the human behavioural experiments, there are often practice trials that might help alleviate the issue and help build up stable criteria rapidly. In addition, a growing number of studies have demonstrated the effects of cross-trial sequence on the human lexical decision performance (e.g., Balota, Aschenbrenner, & Yap, 2016), where stimulus degradation and lexicality in the previous trial have impacted on the responses to the current stimuli, providing evidence for trial-by-trial adjustments to decision making. However, the underlying mechanism remains to be understood. Within the current modelling framework, it is possible to investigate the issue by dynamically adjusting the current stimulus polarity scores with reference to stimulus polarity scores in the previous trial or to implement a more flexible decision mechanism such as the leaky competing accumulator (Usher & McClelland, 2001, 2004).

Another consideration is that the simulations of different types of acquired dyslexia only consider the most typical patients’ lexical decision patterns. Clearly, there are wide variations in reading patterns within each type of dyslexia (Behrmann, et al., 1998; Crisp & Lambon Ralph, 2006; Roberts et al., 2010; Woollams, et al., 2007). The variations could, for instance, result from the severity of reading deficits and premorbid individual differences in reading (e.g., Dilkina, McClelland, & Plaut, 2008). Numerous studies have shown individuals differ with regard to reading experience and vocabulary knowledge as a consequence of great variations in reading effects of skilled readers (Adelman et al., 2012; Andrews & Hersch, 2010; Davies et al., 2017; Yap et al., 2012). Individual differences in the degree of semantic reliance during exception word reading could account for different reading patterns observed in patients with semantic dementia (Woollams et al., 2016). Considering both the severity of reading disorders and premorbid individual differences in simulations of acquired dyslexia would be an interesting topic for further investigation.

## 5. Conclusion

A large-scale computational model of the human visual word recognition system was developed to explore the underlying processing mechanisms in lexical decision. We demonstrated that the model could perform the lexical decision tasks based on the measure of polarity combined across processing layers within the reading system. Importantly, the model was able to account for the graded semantic influences on lexical decision corresponding to various types of foils, providing evidence for semantic access in lexical decision – both in typical and neurologically-impaired reading.

## Acknowledgements

Preliminary data were presented at the 35th annual conference of the Cognitive Science Society. This research was supported by grants under the Cognitive Foresight Initiative (jointly funded by EPSRC, MRC and BBSRC - EP/F03430X/1) and the Neuroscience Research Institute at the University of Manchester. All authors contributed in a significant way to the manuscript. The authors have no conflicts of interest to declare.

## Appendix A: The exploration of control units

In Simulation 1 we asserted that the control units were important for training a network with deep structures, because they allowed the model to learn to manage its own temporal dynamics. This simulation explored this computational issue in more detail.

In the reading process, we expect activation in the network to move progressively forward from the visual areas, and for neurons to remain near their equilibrium states before the forward moving activation reaches them. Evidence for the sequential time course of the letter and word processing in humans comes from several event-related potential (ERP) studies (Grainger et al., 2008; Petit, Midgley, Holcomb, & Grainger, 2006). During word processing, the peak brain activities corresponding to visual, orthographic, phonological and semantic processes occurred at different time periods (Grainger et al., 2008). In the model, however, there is a tendency to develop strong activations in the phonological and semantic units at the very early time ticks, in particular when using the pre-trained weights. This could potentially prevent the model from learning. To overcome this issue, it is possible to use the control units to regulate when individual layers are able to activate. The following sections illustrate how using pre-trained weights can inhibit learning, and how control units can address this problem, by learning to control the temporal dynamics of the network.

### A.1. The Pitfalls of Using Pre-trained Weights

Using pre-trained weights could be helpful in two aspects when training a large-scale recurrent network. First, it can potentially shorten the training time because a large network can generally be dissembled into several small feedforward networks. These small networks can be trained separately and the weights obtained from each part can be embedded into the original large network for global training. Second, the pre-trained weights can be used in an ecologically valid manner to simulate pre-existing knowledge within the network before it starts to learn a new skill - as in Harm and Seidenberg’s (2004) reading model and the recurrent reading model in this paper, where a degree of competency in the phonology-semantics component is developed prior to reading acquisition.

Despite these attractive benefits, there is a complication in using pre-trained weights - if only parts of the network use the pre-trained weights, the rest of the network could become very difficult to train, in particular where there are units which are connected partly through pre-trained weights and partly through new untrained weights. The major inputs to the units would come from the pre-trained weights side because the connection values of the pre-trained weights are generally rather large in comparison with the untrained weights. This potentially weakens the ability of the network to learn through error propagation. To take an example, considering a simple two-layer feedforward network, it has connections as shown in Figure A1. A unit *k* is connected to a unit *i* through *w_i_*, a frozen, pre-trained weight value of 10 and to a unit *j* through *w_j_*, with a random initial weight value of 0.1. If both unit *i* and unit *j* have the output values of 1, the output of unit *k* can be written as:

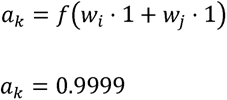

where *f*(·) is a logistic function. If the desired output of the unit *k*, *t_k_* =1 and the *η* = 0.1, according to the gradient decent the weight change Δ*w_j_* for unit *j* is

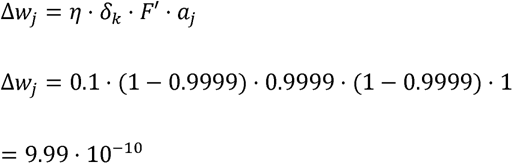

where *δ_k_* = (*t_k_* − *a_k_*) and *F*’ = *a_k_* · (1 − *a_k_*),

The weight change for unit *i* is zero because the weights are frozen. Although the weight change is quite small, this might be satisfied because the output of the unit *k* is very close to the desired output so that the extra training may not be necessary. However, if the desired output *t_k_* = −1, according to the same formula, the weight change Δ*w_j_* is still only Δ*w_j_* = −1.9999 · 10^−5^. Thus, it may result in extremely long (or failed) training because the weight changes are tiny due to the use the frozen pre-trained weights.

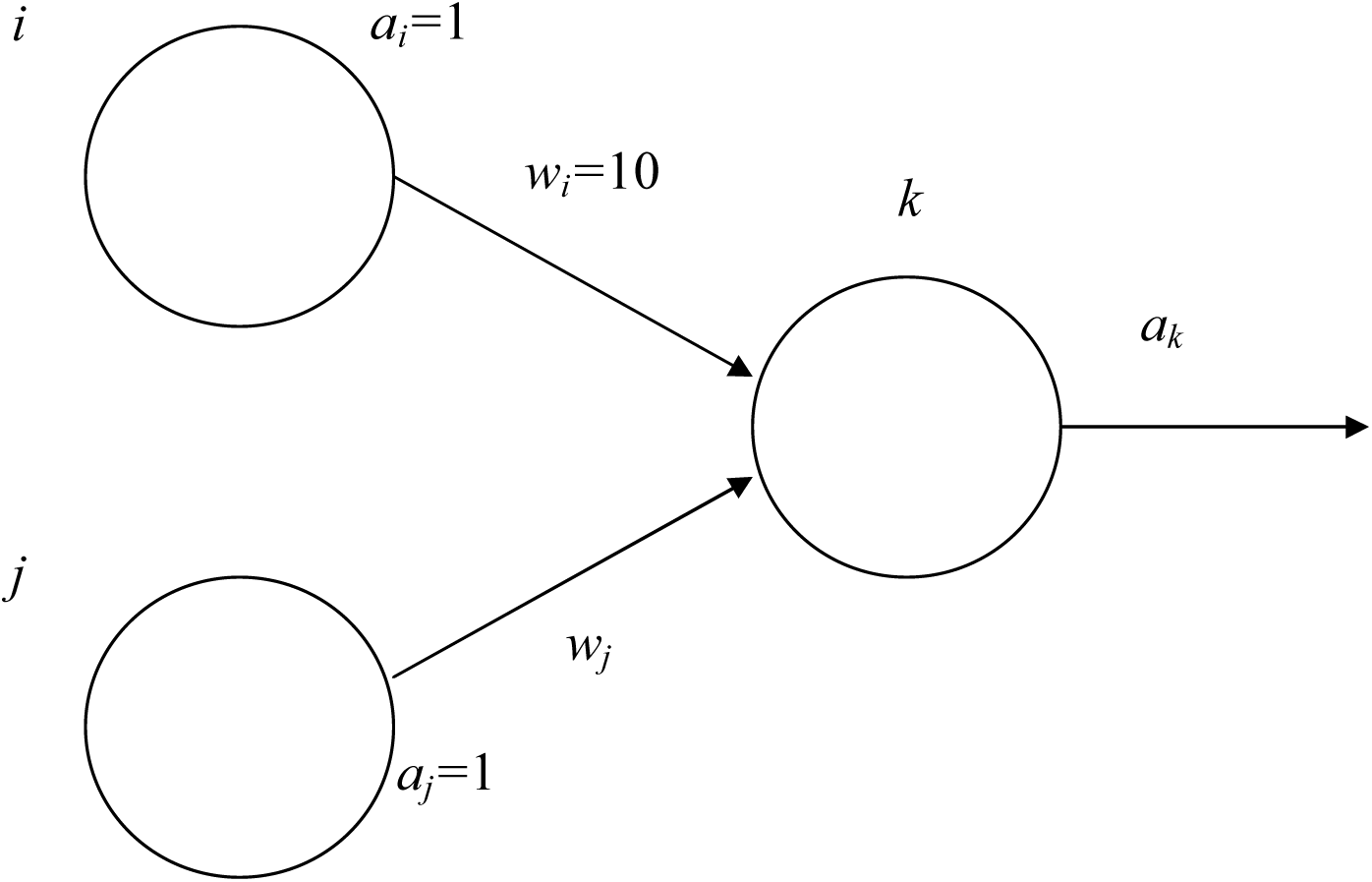

### A.2. The Use of Control Units

One way of resolving both of these issues is through control units (Dilkina et al., 2010), which provide a mechanism to regulate the temporal dynamics of activation within the model including, but not limited to the contribution from units attached to pre-trained weights. The control technique can be implemented by using a control unit along with a strong negative weight connected to every unit it needs to control. The control unit will be free to learn to regulate the activity of the units it controls in whatever fashion reduces errors, but we anticipate that for control units connected to other units with pre-trained weights it will learn to exert a strong inhibition early in training which will gradually decrease as the non pre-trained weights start to participate in the task. In the case of the network illustrated in Figure A1, the control units can be connected to the units *i* and *j*, and might initially learn to reduce the input contribution from unit *i* whilst preserving it from unit *k*. Similarly, for units at deep layers within a large dynamic network, we anticipate that the control units would act to inhibit activation during the first few time ticks when useful information has not yet had time to accumulate at the deeper levels of the network.

The results in Simulation 1 (Figure 3) clearly demonstrate the performance benefits of using control units, in particular for semantics. To examine how they have achieved this, we computed the inhibitory effects of the control units on the H1, H2, H3 and H4 layers (see Figure 1). Figure A2 shows the average inhibition effects on these layers as a function of time ticks. For the units that control the deeper layers with pre-trained weights (H3 and H4), the pattern is as expected with a high initial level of inhibition that decreases towards zero with time. This is in contrast with H2, which starts with very low levels of inhibition and maintains this throughout processing. Presumably reflecting the importance of allowing information to progress from the visual layers to the phonological layers as quickly as possible. One might have expected the same pattern for H1, which controls the information flow along the semantic route, but here we see high initial levels of inhibition that decline only slightly throughout the whole course of processing. This suggests that activation of the semantic system is achieved mainly through the indirect route (via phonology). Overall, these analyses of the time dynamics of control units in the model confirm that the control units improve the performance of the model by regulating the temporal dynamics of each layer in the model to allow efficient processing.

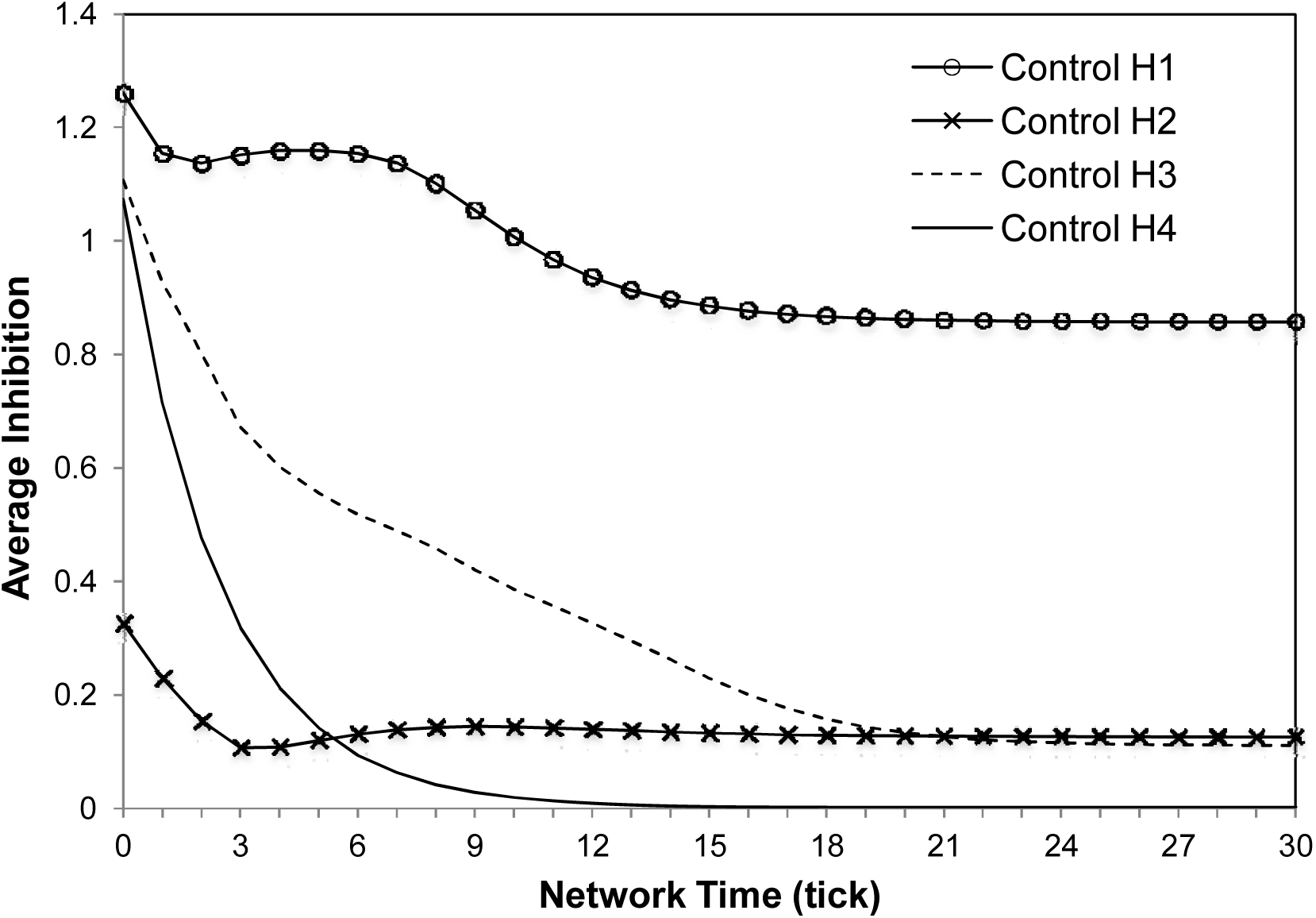

## Footnotes

1 Note this vowel centred alignment is not necessary for good performance in the model. Chang et al. (2012a) showed that any reasonable approximation of the optimal viewing position (OVP) with a central position slightly to the left of centre would produce similar results.

## References

Adelman, J. S., Sabatos-DeVito, M. G., Marquis, S. J., & Estes, Z. (2014). Individual differences in reading aloud: A mega-study, item effects, and some models. Cognitive Psychology, 68, 113–160. doi: 10.1016/j.cogpsych.2013.11.001

Andrews, S. (1982). Phonological recoding - Is the regularity effect consistent. Memory & Cognition, 10(6), 565–575. doi: 10.3758/bf03202439

Andrews, S. (1989). Frequency and neighborhood effects on lexical access - Activation or search. Journal of Experimental Psychology-Learning Memory and Cognition, 15(5), 802–814. doi: 10.1037/0278-7393.15.5.802

Andrews, S. (1992). Frequency and neighborhood effects on lexical access - Lexical similarity or orthographic redundancy. Journal of Experimental Psychology-Learning Memory and Cognition, 18(2), 234–254. doi: 10.1037/0278-7393.18.2.234

Andrews, S. (1997). The effect of orthographic similarity on lexical retrieval: Resolving neighborhood conflicts. Psychonomic bulletin & review, 4(4), 439–461. doi: 10.3758/BF03214334

Andrews, S., & Hersch, J. (2010). Lexical precision in skilled readers: Individual differences in masked neighbor priming. J Exp Psychol Gen, 139(2), 299–318. doi: 10.1037/a0018366

Arguin, M., Fiset, S., & Bub, D. (2002). Sequential and parallel letter processing in letter-by-letter dyslexia. Cognitive Neuropsychology, 19(6), 535–555. doi: 10.1080/02643290244000040.

Azuma, T., & Van Orden, G. C. (1997). Why SAFE is better than FAST: The relatedness of a word’s meanings affects lexical decision times. Journal of Memory and Language, 36(4), 484–504. doi: 10.1006/jmla.1997.2502

Baron, J., & Strawson, C. (1976). Use of orthographic and word specific knowledge in reading words aloud. Journal of Experimental Psychology-Human Perception and Performance, 2(3), 386–393. doi: 10.1037/0096-1523.2.3.386

Balota DA, Aschenbrenner AJ, & Yap MJ (2016). Dynamic adjustment of lexical processing in the lexical decision task: Cross-trial sequence effects. The Quarterly Journal of Experimental Psychology:1–10. doi:10.1080/17470218.2016.1240814

Balota, D. A., & Chumbley, J. I. (1984). Are lexical decisions a good measure of lexical access - the role of word-frequency in the neglected decision stage. Journal of Experimental Psychology-Human Perception and Performance, 10(3), 340–357. doi: 10.1037/0096-1523.10.3.340

Balota, D. A., Cortese, M. J., Sergent-Marshall, S. D., Spieler, D. H. & Yap, M. J. (2004). Visual word recognition of single-syllable words. Journal of Experimental Psychology-General, 133(2), 283–316. doi:10.1037/0096-3445.133.2.283

Balota, D. A., Yap, M. J., Hutchison, K. A., Cortese, M. J., Kessler, B., Loftis, B., Treiman, R. (2007). The English Lexicon Project. Behavior Research Methods, 39(3), 445–459. doi: 10.3758/bf03193014

Beauvois, M. F., & Derouesne, J. (1979). Phonological alexia - 3 dissociations. Journal of Neurology Neurosurgery and Psychiatry, 42(12), 1115–1124. doi: 10.1136/jnnp.42.12.1115

Becker, C. A. (1980). Semantic context effects in visual word recognition: An analysis of semantic strategies. Memory & Cognition, 8(6), 493–512. doi: 10.3758/BF03213769

Behrmann, M., Plaut, D. C., & Nelson, J. (1998). A literature review and new data supporting an interactive account of letter-by-letter reading. Cognitive Neuropsychology, 15(1-2), 7–51. doi: 10.1080/026432998381212

Benedet, M., Patterson, K., Gomez-Pastor, I., & de la Rocha, M. L. G. (2006). “Non-Semantic” aspects of language in semantic dementia: As normal as they’re said to be? Neurocase, 12(1), 15–26. doi: 10.1080/13554790500446868

Binder, J. R., Desai, R. H., Graves, W. W., & Conant, L. L. (2009). Where is the semantic system? A critical review and meta-analysis of 120 functional neuroimaging studies. Cerebral Cortex, 19(12), 2767–2796. doi: 10.1093/cercor/bhp055

Binder, J. R., Frost, J. A., Hammeke, T. A., Bellgowan, P. S. F., Rao, S. M., & Cox, R. W. (1999). Conceptual processing during the conscious resting state: a functional mri study. Journal of Cognitive Neuroscience, 11(1), 80–93. doi: 10.1162/089892999563265

Binder, J. R., McKiernan, K. A., Parsons, M. E., Westbury, C. F., Possing, E. T., Kaufman, J. N., & Buchanan, L. (2003). Neural correlates of lexical access during visual word recognition. Journal of Cognitive Neuroscience, 15(3), 372–393. doi: 10.1162/089892903321593108

Bishop, C. M. (2006). Pattern Recognition and Machine Learning (Vol. 4): Springer New York.

Blazely, A. M., Coltheart, M., & Casey, B. J. (2005). Semantic impairment with and without surface dyslexia: Implications for models of reading. Cognitive Neuropsychology, 22(6), 695–717. doi: 10.1080/02643290442000257

Borowsky, R., & Besner, D. (1993). Visual word recognition - a multistage activation model. Journal of Experimental Psychology-Learning Memory and Cognition, 19(4), 813–840. doi: 10.1037/0278-7393.19.4.813

Borowsky, R., & Besner, D. (2006). Parallel distributed processing and lexical-semantic effects in visual word recognition: are a few stages necessary? Psychol Rev, 113(1), 181–195; discussion 196-200. doi: 10.1037/0033-295x.113.1.181

Borowsky, R., & Masson, M. E. J. (1996). Semantic ambiguity effects in word identification. Journal of Experimental Psychology: Learning, Memory, and Cognition, 22(1), 63–85. doi: 10.1037/0278-7393.22.1.63

Bonner, M. F., Vesely, L., Price, C., Anderson, C., Richmond, L., Farag, C., Avants, & Grossman, M. (2009). Reversal of the concreteness effect in semantic dementia. Cognitive Neuropsychology, 26(6), 568–579. doi: 10.1080/02643290903512305

Breedin, S. D., Saffran, E. M., & Coslett, H. B. (1994). Reversal of the concreteness effect in a patient with semantic dementia. Cognitive Neuropsychology, 11(6), 617–660. doi: 10.1080/02643299408251987

Busemeyer, J. R., & Townsend, J. T. (1993). Decision field theory: a dynamic-cognitive approach to decision making in an uncertain environment. Psychological Review, 100(3), 432–459. doi: 10.1037/0033-295X.100.3.432

Caccappolo-van Vliet, E., Miozzo, M., & Stern, Y. (2004). Phonological dyslexia without phonological impairment? Cognitive Neuropsychology, 21(8), 820–839. doi: 10.1080/02643290342000465

Chang, Y. N., Furber, S., & Welbourne, S. (2012a). “Serial” effects in parallel models of reading. Cognitive Psychology, 64(4), 267–291. doi: 10.1016/j.cogpsych.2012.01.002

Chang, Y. N., Furber, S., & Welbourne, S. (2012b). Generating realistic codes for use in neural network models. In N. Miyake, D. Peebles, & R.P. Cooper (Eds.), Proceedings of the 34th Annual Confrerence of the Cognitive Science Society (pp. 198–203). Austin, Tx: Cognitive Science Society.

Chang, Y.-N., & Monaghan, P. (2018). Quantity and Diversity of Preliteracy Language Exposure Both Affect Literacy Development: Evidence from a Computational Model of Reading. Scientific Studies of Reading, 1–19. doi:10.1080/10888438.2018.1529177

Chang, Y.-N., Monaghan, P., & Welbourne, S. (2019). A computational model of reading across development: Effects of literacy onset on language processing. Journal of Memory and Language, 108, 104025. doi: https://doi.org/10.1016/j.jml.2019.05.003

Cheyette, S. J., & Plaut, D. C. (2017). Modeling the N400 ERP component as transient semantic over-activation within a neural network model of word comprehension. Cognition, 162, 153–166. doi: https://doi.org/10.1016/j.cognition.2016.10.016

Chumbley, J. I., & Balota, D. A. (1984). A word’s meaning affects the decision in lexical decision. Memory & Cognition, 12(6), 590–606. doi: 10.3758/BF03213348

Collins, A. M., & Loftus, E. F. (1975). A spreading-activation theory of semantic processing. Psychological Review, 82(6), 407–428. doi: 10.1037/0033-295X.82.6.407

Coltheart, M., Besner, D., Jonasson, J. T., & Davelaar, E. (1979). Phonological encoding in the lexical decision task. Quarterly Journal of Experimental Psychology, 31(3), 489–507. doi: 10.1080/14640747908400741

Coltheart, M., Davelaar, E., Jonasson, J. T., & Besner, D. (1977). Access to the internal lexicon. In S. Dornic (Ed.), Attention and Performance VI (pp. 535–555). Hillsdale, NJ: Lawrence Erlbaum Associates.

Coltheart, M. (2004). Are there lexicons? The Quarterly Journal of Experimental Psychology Section A, 57(7), 1153–1171. doi: 10.1080/02724980443000007

Coltheart, M., Rastle, K., Perry, C., Langdon, R., & Ziegler, J. (2001). DRC: A dual route cascaded model of visual word recognition and reading aloud. Psychological Review, 108(1), 204–256. doi: 10.1037//0033-295x.108.1.204

Cortese, M. J., & Khanna, M. M. (2008). Age of acquisition ratings for 3,000 monosyllabic words. Behavior Research Methods, 40(3), 791–794. doi: 10.3758/brm.40.3.791

Coslett, H. B., & Saffran, E. M. (1989). Evidence for preserved reading in pure alexia. Brain, 112, 327–359. doi: 10.1093/brain/112.2.327

Coslett, H. B., & Saffran, E. M. (1994). Mechanisms of implicit reading in alexia. In M. J. Farah & G. Ratcliff (Eds.), The Neuropsychology of High-Level Vision (pp. 299–330). Hillsdale, NJ: Lawrence Erlbaum Associates Inc.

Crisp, J., & Lambon Ralph, M. A. (2006). Unlocking the nature of the phonological–deep dyslexia continuum: The keys to reading aloud are in phonology and semantics. Journal of Cognitive Neuroscience, 18(3), 348–362. doi: 10.1162/jocn.2006.18.3.348

Cuetos, F., ValleArroyo, F., & Suarez, M. P. (1996). A case of phonological dyslexia in spanish. Cognitive Neuropsychology, 13(1), 1–24. doi: 10.1080/026432996382042

Cushman, C. L., & Johnson, R. L. (2011). Age-of-Acquisition effects in pure alexia. The Quarterly Journal of Experimental Psychology, 64(9), 1726–1742. doi: 10.1080/17470218.2011.556255

Damasio, A. R., & Damasio, H. (1983). The anatomic basis of pure alexia. Neurology, 33(12), 1573–1583. doi: 10.1212/WNL.33.12.1573

Davies, R. A. I., Arnell, R., Birchenough, J. M. H., Grimmond, D., & Houlson, S. (2017). Reading through the life span: Individual differences in psycholinguistic effects. J Exp Psychol Learn Mem Cogn, 43(8), 1298–1338. doi: 10.1037/xlm0000366

Degroot, A. M. B. (1989). Representational aspects of word imageability and word-frequency as assessed through word-association. Journal of Experimental Psychology-Learning Memory and Cognition, 15(5), 824–845. doi: 10.1037/0278-7393.15.5.824

Denes, G., Cipolotti, L., & Semenza, C. (1987). How does a phonological dyslexic read words she has never seen. Cognitive Neuropsychology, 4(1), 11–31. doi: 10.1080/02643298708252032

Diesfeldt, H. F. A. (1992). Impaired and preserved semantic memory functions in dementia. In B. Lars (Ed.), Advances in Psychology (Vol. 89, pp. 227–263). Oxford, England: North-Holland.

Dilkina, K., McClelland, J. L., & Plaut, D. C. (2008). A single-system account of semantic and lexical deficits in five semantic dementia patients. Cognitive Neuropsychology, 25(2), 136–164. doi: 10.1080/02643290701723948

Dilkina, K., McClelland, J. L., & Plaut, D. C. (2010). Are there mental lexicons? The role of semantics in lexical decision. Brain Research, 1365, 66–81. doi: 10.1016/j.brainres.2010.09.057

Dujardin, T., Etienne, Y., Contentin, C., Bernard, C., Largy, P., Mellier, D., Lalonde, R., & Rebai, M. (2011). Behavioral performances in participants with phonological dyslexia and different patterns on the N170 component. Brain and Cognition, 75(2), 91–100. doi: 10.1016/j.bandc.2010.10.006

Evans, G. A. L., Lambon Ralph, M. A., & Woollams, A. M. (2012). What’s in a word? A parametric study of semantic influences on visual word recognition. Psychonomic Bulletin & Review, 19(2), 325–331. doi: 10.3758/s13423-011-0213-7

Fiset, D., Arguin, M., & McCabe, E. (2006). The breakdown of parallel letter processing in letter-by-letter dyslexia. Cognitive Neuropsychology, 23(2), 240–260. doi: 10.1080/02643290442000437

Forster, K. I. (1976). Accessing the mental lexicon. In R.J. Wales & E. Walker (Eds.), New Approaches to Language Mechanisms. (pp. 257–287). Amsterdam: North-Holland

Friedman, R. B., & Hadley, J. A. (1992). Letter-by-letter surface alexia. Cognitive Neuropsychology, 9(3), 185–208. doi: 10.1080/02643299208252058

Fujimaki, N., Hayakawa, T., Ihara, A., Wei, Q., Munetsuna, S., Terazono, Y., Matani, A., & Murata, T. (2009). Early neural activation for lexico-semantic access in the left anterior temporal area analyzed by an fMRI-assisted MEG multidipole method. Neuroimage, 44(3), 1093–1102. doi: 10.1016/j.neuroimage.2008.10.021

Garrard, P., Lambon Ralph, M. A., Hodges, J. R., & Patterson, K. (2001). Prototypicality, distinctiveness, and intercorrelation: Analyses of the semantic attributes of living and nonliving concepts. Cognitive Neuropsychology, 18(2), 125–174. doi: 10.1080/02643290125857

Grainger, J., & Jacobs, A. M. (1996). Orthographic processing in visual word recognition: A multiple read-out model. Psychological Review, 103(3), 518–565. doi: 10.1037/0033-295x.103.3.518

Grainger, J., Muneaux, M., Farioli, F., & Ziegler, J. C. (2005). Effects of phonological and orthographic neighbourhood density interact in visual word recognition. The Quarterly Journal of Experimental Psychology Section A, 58(6), 981–998. doi:10.1080/02724980443000386

Grondin, R., Lupker, S. J., & McRae, K. (2009). Shared Features Dominate Semantic Richness Effects for Concrete Concepts. Journal of Memory and Language, 60(1), 1–19. doi: 10.1016/j.jml.2008.09.001

Harm, M. W., & Seidenberg, M. S. (1999). Phonology, reading acquisition, and dyslexia: insights from connectionist models. Psychological Review, 106(3), 491–528. doi: 10.1037/0033-295X.106.3.491

Harm, M. W., & Seidenberg, M. S. (2004). Computing the meanings of words in reading: cooperative division of labor between visual and phonological processes. Psychological Review, 111(3), 662–720. doi: 10.1037/0033-295x.111.3.662

Hauk, O., Coutout, C., Holden, A., & Chen, Y. (2012). The time-course of single-word reading: evidence from fast behavioral and brain responses. Neuroimage, 60(2), 1462–1477. doi: 10.1016/j.neuroimage.2012.01.061

Hauk, O., Patterson, K., Woollams, A., Watling, L., Pulvermuller, F., & Rogers, T. T. (2006). Q: When would you prefer a sossage to a sausage? A: At about 100 msec. ERP correlates of orthographic typicality and lexicality in written word recognition. Journal of Cognitive Neuroscience, 18(5), 818–832. doi: 10.1162/jocn.2006.18.5.818

Hino, Y., & Lupker, S. J. (1996). Effects of polysemy in lexical decision and naming: An alternative to lexical access accounts. Journal of Experimental Psychology: Human Perception and Performance, 22(6), 1331. doi: 10.1037/0096-1523.22.6.1331

Hodges, J. R., Patterson, K., Oxbury, S., & Funnell, E. (1992). Semantic dementia - Progressive fluent aphasia with temporal-lobe atrophy. Brain, 115, 1783–1806. doi: 10.1093/brain/115.6.1783

Hoffman, P., & Lambon Ralph, M. A. (2011). Reverse Concreteness Effects Are Not a Typical Feature of Semantic Dementia: Evidence for the Hub-and-Spoke Model of Conceptual Representation. Cerebral Cortex, 21(9), 2103–2112. doi: 10.1093/cercor/bhq288

Hoffman, P., Lambon Ralph, M. A., & Woollams, A. M. (2015). Triangulation of the neurocomputational architecture underpinning reading aloud. Proceedings of the National Academy of Sciences of the United States of America - PNAS, 112(28), E3719–E3728. doi: 10.1073/pnas.1502032112

Hoffman, P., Jones, R. W., & Lambon Ralph, M. A. (2013). Be concrete to be comprehended: Consistent imageability effects in semantic dementia for nouns, verbs, synonyms and associates. Cortex, 49(5), 1206–1218. doi: 10.1016/j.cortex.2012.05.007

Hoffman, P., & Woollams, A. M. (2015). Opposing effects of semantic diversity in lexical and semantic relatedness decisions. J Exp Psychol Hum Percept Perform, 41(2), 385–402. doi: 10.1037/a0038995

Howard, D. (1991). Letter-by-letter readers: Evidence for parallel processing. Hillsdale, NJ, US: Lawrence Erlbaum Associates, Inc.

James, C. T. (1975). Role of semantic information in lexical decisions. Journal of Experimental Psychology-Human Perception and Performance, 104(2), 130–136. doi: 10.1037/0096-1523.1.2.130

Jared, D., McRae, K., & Seidenberg, M. S. (1990). The basis of consistency effects in word naming. Journal of Memory and Language, 29(6), 687–715. doi: 10.1016/0749-596X(90)90044-Z

Jastrzembski, J. E. (1981). Multiple meanings, number of related meanings, frequency of occurrence, and the lexicon. Cognitive Psychology, 13(2), 278–305. doi: 10.1016/0010-0285(81)90011-6

Jefferies, E., Patterson, K., Jones, R. W., & Lambon Ralph, M. A. (2009). Comprehension of concrete and abstract words in semantic dementia. Neuropsychology, 23(4), 492–499. doi: 10.1037/a0015452

Jefferies, E., Lambon Ralph, M. A., Jones, R., Bateman, D., & Patterson, K. (2004). Surface dyslexia in semantic dementia: a comparison of the influence of consistency and regularity. Neurocase, 10(4), 290–299. doi: 10.1080/13554790490507623

Johnson, R. L., & Rayner, K. (2007). Top-down and bottom-up effects in pure alexia: Evidence from eye movements. Neuropsychologia, 45(10), 2246–2257. doi: 10.1016/j.neuropsychologia.2007.02.026

Joordens, S., & Becker, S. (1997). The long and short of semantic priming effects in lexical decision. Journal of Experimental Psychology-Learning Memory and Cognition, 23(5), 1083–1105. doi: 10.1037/0278-7393.23.5.1083

Kello, C. T. (2006). Considering the junction model of lexical processing. In S. Andrews (Ed.), From inkmarks to ideas: Current issues in lexical processing (pp. 50–75). New York: Psychology Press.

Kellas, G., Ferraro, F. R., & Simpson, G. B. (1988). Lexical ambiguity and the timecourse of attentional allocation in word recognition. Journal of Experimental Psychology: Human Perception and Performance, 14(4), 601. doi: 10.1037/0096-1523.14.4.601

Kroll, J. F., & Merves, J. S. (1986). Lexical access for concrete and abstract words. Journal of Experimental Psychology-Learning Memory and Cognition, 12(1), 92–107. doi: 10.1037/0278-7393.12.1.92

Landauer, T. K., Foltz, P. W., & Laham, D. (1998). An introduction to latent semantic analysis. Discourse Processes, 25(2-3), 259–284. doi: 10.1080/01638539809545028

Laszlo, S., & Plaut, D. C. (2012). A neurally plausible parallel distributed processing model of event-related potential word reading data. Brain and Language, 120(3), 271–281. doi: 10.1016/j.bandl.2011.09.001

Lund, K., & Burgess, C. (1996). Producing high-dimensional semantic spaces from lexical co-occurrence. Behavior Research Methods Instruments & Computers, 28(2), 203–208. doi: 10.3758/BF03204766

Lupker, S. J., & Pexman, P. M. (2010). Making things difficult in lexical decision: The impact of pseudohomophones and transposed-letter nonwords on frequency and semantic priming effects. Journal of Experimental Psychology-Learning Memory and Cognition, 36(5), 1267–1289. doi: 10.1037/a0020125

Martensen, H., Dijkstra, T., & Maris, E. (2005). A werd is not quite a word: On the role of sublexical phonological information in visual lexical decision. Language and Cognitive Processes, 20(4), 513–552. doi:10.1080/01690960444000043

McClelland, J. L., & Rumelhart, D. E. (1981). An interactive activation model of context effects in letter perception: Part 1. An account of basic findings. Psychological Review, 88(5), 375–407.

McRae, K., Cree, G. S., Seidenberg, M. S., & McNorgan, C. (2005). Semantic feature production norms for a large set of living and nonliving things. Behavior Research Methods, 37(4), 547–559. doi: 10.3758/BF03192726

Meyer, D., & Schvaneveldt, R. (1971). Facilitation in recognizing pairs of words: Evidence of a dependence between retrieval operations. Journal of Experimental Psychology, 90(2), 227–234. doi: 10.1037/h0031564

Meyer, D. E., Schvaneveldt, R. W., & Ruddy, M. G. (1974). Functions of graphemic and phonemic codes in visual word-recognition. Memory & Cognition, 2(2), 309–321. doi: 10.3758/bf03209002

Miller, G. A., Beckwith, R., Fellbaum, C., Gross, D., & Miller, K. J. (1990). Introduction to wordnet: An on-line lexical database. International Journal of Lexicography, 3(4), 235–244. doi: 10.1093/ijl/3.4.235

Millis, M., & Bution, S. (1989). The effect of polysemy on lexical decision time: Now you see it, now you don’t. Memory & Cognition, 17(2), 141–147. doi: 10.3758/BF03197064

Milota, V. C., Widau, A. A., McMickell, M. R., Juola, J. F., & Simpson, G. B. (1997). Strategic reliance on phonological mediation in lexical access. Memory & Cognition, 25(3), 333–344. doi: 10.3758/bf03211289

Monaghan, P., Chang, Y. N., Welbourne, S., & Brysbaert, M. (2017). Exploring the relations between word frequency, language exposure, and bilingualism in a computational model of reading. Journal of Memory and Language, 93, 1–21. doi: 10.1016/j.jml.2016.08.003

Monaghan, P., Shillcock, R., & McDonald, S. (2004). Hemispheric asymmetries in the split-fovea model of semantic processing. Brain and Language, 88(3), 339–354

Montant, M., & Behrmann, M. (2001). Phonological activation in pure alexia. Cognitive Neuropsychology, 18(8), 697–727. doi: 10.1080/02643290143000042

Morton, J. (1969). Interaction of information in word recognition. Psychological Review, 76(2), 165–178. doi: 10.1037/h0027366

Norris, D. (2006). The bayesian reader: explaining word recognition as an optimal bayesian decision process. Psychological Review, 113(2), 327–357. doi: 10.1037/0033-295x.113.2.327

Norris, D. (2009). Putting it all together: A unified account of word recognition and reaction-time distributions. Psychological Review, 116(1), 207–219. doi: 10.1037/a0014259

Parkin, A. J., & Underwood, G. (1983). Orthographic vs. phonological irregularity in lexical decision. Memory & Cognition, 11(4), 351–355. doi: 10.3758/BF03202449

Patterson, K. (1982). The Relation between Reading and Phonological Coding: Further Neuropsychological Observations. In A. W. Ellis (Ed.), Normality and Pathology in Cognitive Functions (pp. 77–111). London, UK: Academic Press.

Patterson, K., & Kay, J. (1982). Letter-by-letter reading: Psychological descriptions of a neurologial syndrome. The Quarterly Journal of Experimental Psychology Section A, 34(3), 411–441. doi: 10.1080/14640748208400852

Patterson, K., & Marcel, A. (1992). Phonological alexia or phonological alexia? In D. Alegria, D. Holender, J. Junca de Morais, & M. Radeau (Eds), Analytic Approaches to Human Cognition (pp. 259–274). Oxford, England: North-Holland.

Patterson, K., Nestor, P. J., & Rogers, T. T. (2007). Where do you know what you know? The Representation of semantic knowledge in the human brain. Nature Reviews Neuroscience, 8(12), 976–987. doi: 10.1038/nrn2277

Patterson, K., Lambon Ralph, M. A., Jefferies, E., Woollams, A., Jones, R., Hodges, J. R., & Rogers, T. T. (2006). “Presemantic” cognition in semantic dementia: Six deficits in search of an explanation. Journal of Cognitive Neuroscience, 18(2), 169–183. doi: 10.1162/089892906775783714

Patterson, K. E., & Marcel, A. J. (1977). Aphasia, dyslexia and phonological coding of written words. Quarterly Journal of Experimental Psychology, 29 (May), 307–318. doi: 10.1080/14640747708400606

Pearlmutter, B. A. (1989). Learning state space trajectories in recurrent neural networks. Neural Computation, 1(2), 263–269. doi:10.1162/neco.1989.1.2.263

Pearlmutter, B. A. (1995). Gradient calculations for dynamic recurrent neural networks – a survey. IEEE Transactions on Neural Networks, 6(5), 1212–1228. doi: 10.1109/72.410363

Perry, C., Ziegler, J. C., & Zorzi, M. (2007). Nested incremental modeling in the development of computational theories: The CDP+ model of reading aloud. Psychological Review, 114(2), 273–315. doi: 10.1037/0033-295x.114.2.273

Petit, J. P., Midgley, K. J., Holcomb, P. J., & Grainger, J. (2006). On the time course of letter perception: a masked priming erp investigation. Psychonomic Bulletin & Review, 13(4), 674–681. doi: 10.3758/BF03193980

Pineda, F. J. (1987). Generalization of back-propagation to recurrent neural networks. Physical Review Letters, 59(19), 2229–2232. doi: 10.1103/PhysRevLett.59.2229

Plaut, D. C. (1997). Structure and function in the lexical system: insights from distributed models of word reading and lexical decision. Language and Cognitive Processes, 12(5-6), 765–805. doi: 10.1080/016909697386682

Plaut, D. C., & Booth, J. R. (2006). More modeling but still no stages: Reply to Borowsky and Besner. Psychological Review, 113(1), 196–200.

Plaut, D. C., McClelland, J. L., Seidenberg, M. S., & Patterson, K. (1996). Understanding normal and impaired word reading: computational principles in quasi-regular domains. Psychological Review, 103(1), 56–115. doi: 10.1037/0033-295X.103.1.56

Plaut, D. C., & Shallice, T. (1993). Deep dyslexia: A case study of connectionist neuropsychology. Cognitive Neuropsychology, 10(5), 377–500. doi: 10.1080/02643299308253469

Rabovsky, M., & McRae, K. (2014). Simulating the N400 ERP component as semantic network error: Insights from a feature-based connectionist attractor model of word meaning. Cognition, 132 (1), 68–89. doi: 10.1016/j.cognition.2014.03.010

Rastle, K., Harrington, J., & Coltheart, M. (2002). 358,534 nonwords: The ARC Nonword Database. The Quarterly Journal of Experimental Psychology Section A, 55(4), 1339–1362. doi: 10.1080/02724980244000099

Ratcliff, R., Gomez, P., & McKoon, G. (2004). A diffusion model account of the lexical decision task. Psychological Review, 111(1), 159–182. doi: 10.1037/0033-295x.111.1.159

Rapcsak, S. Z., Beeson, P. M., Henry, M. L., Leyden, A., Kim, E., Rising, K., Andersen, S., Cho, H. (2009). Phonological dyslexia and dysgraphia: Cognitive mechanisms and neural substrates. Cortex, 45(5), 575–591. doi:/10.1016/j.cortex.2008.04.006

Reilly, J., Grossman, M., & McCawley, G. (2006). Concreteness effects in lexical processing of semantic dementia. Brain and Language, 99(1), 157–158. doi: https://doi.org/10.1016/j.bandl.2006.06.088

Roberts, D. J., Lambon Ralph, M. A., & Woollams, A. M. (2010). When does less yield more? The impact of severity upon implicit recognition in pure alexia. Neuropsychologia, 48(9), 2437–2446. doi: 10.1016/j.neuropsychologia.2010.04.002

Roder, B., Kusmierek, A., Spence, C., & Schicke, T. (2007). Developmental vision determines the reference frame for the multisensory control of action. Proceedings of the National Academy of Sciences of the United States of America, 104(11), 4753–4758. doi: 10.1073/pnas.0607158104

Rogers, T. T., Lambon Ralph, M. A., Garrard, P., Bozeat, S., McClelland, J. L., Hodges, J. R., & Patterson, K. (2004). Structure and deterioration of semantic memory: A neuropsychological and computational investigation. Psychological Review, 111(1), 205–235. doi: 10.1037/0033-295x.111.1.205

Rogers, T. T., Lambon Ralph, M. A., Hodges, J. R., & Patterson, K. (2004). Natural selection: The impact of semantic impairment on lexical and object decision. Cognitive Neuropsychology, 21(2-4), 331–352. doi: 10.1080/02643290342000366

Rohde, D. L. T., Gonnerman, L., & Plaut, D. C. (2006). An improved model of semantic similarity based on lexical co-occurrence. Communications of the ACM, 8, 627–633. doi: 10.1.1.131.9401

Rubenstein, H., Garfield, L., & Millikan, J. A. (1970). Homographic entries in the internal lexicon. Journal of Verbal Learning and Verbal Behavior, 9(5), 487–494. doi: 10.1016/S0022-5371(70)80091-3

Rubenstein, H., Lewis, S. S., & Rubenstein, M. A. (1971). Evidence for phonemic recording in visual word recognition. Journal of Verbal Learning and Verbal Behavior, 10(6), 645–657. doi: 10.1016/s0022-5371(71)80071-3

Seidenberg, M. S., & McClelland, J. L. (1989). A distributed, developmental model of word recognition and naming. Psychological Review, 96(4), 523–568. doi: 10.1037/0033-295X.96.4.523

Seidenberg, M. S., & McClelland, J. L. (1990). More words but still no lexicon - Reply. Psychological Review, 97(3), 447–452. doi: 10.1037/0033-295x.97.3.447

Seidenberg, M. S., Waters, G. S., Barnes, M. A., & Tanenhaus, M. K. (1984). When does irregular spelling or pronunciation influence word recognition? Journal of Verbal Learning and Verbal Behavior, 23(3), 383–404. doi: 10.1016/s0022-5371(84)90270-6

Shulman, H. G., & Davison, T. C. B. (1977). Control properties of semantic coding in a lexical decision task. Journal of Verbal Learning and Verbal Behavior, 16(1), 91–98. doi: 10.1016/s0022-5371(77)80010-8

Spieler, D. H., & Balota, D. A. (1997). Bringing Computational Models of Word Naming Down to the Item Level. Psychological Science, 8(6), 411–416. doi: doi:10.1111/j.1467-9280.1997.tb00453.x

Stanovich, K. E., & Bauer, D. W. (1978). Experiments on spelling-to-sound regularity effect in word recognition. Memory & Cognition, 6(4), 410–415. doi: 10.3758/BF03197473

Stone, G. O., & Van Orden, G. C. (1993). Strategic control of processing in word recognition. Journal of Experimental Psychology: Human Perception and Performance, 19(4), 744–774. doi: 10.1037/0096-1523.19.4.744

Taraban, R., & McClelland, J. L. (1987). Conspiracy effects in word pronunciation. Journal of Memory and Language, 26(6), 608–631. doi: 10.1016/0749-596X(87)90105-7

Tree, J. J., & Kay, J. (2006). Phonological dyslexia and phonological impairment: An exception to the rule? Neuropsychologia, 44(14), 2861–2873. doi: 10.1016/j.neuropsychologia.2006.06.006

Usher, M., & McClelland, J. L. (2001). The time course of perceptual choice: The leaky, competing accumulator model. Psychological Review, 108(3), 550–592. doi: 10.1037/0033-295X.108.3.550

Usher, M., & McClelland, J. L. (2004). Loss Aversion and Inhibition in Dynamical Models of Multialternative Choice. Psychological Review, 111(3), 757–769. doi: 10.1037/0033-295X.111.3.757

Vandenberghe, R., Price, C., Wise, R., Josephs, O., & Frackowiak, R. S. J. (1996). Functional anatomy of a common semantic system for words and pictures. Nature, 383(6597), 254–256. doi: 10.1038/383254a0

Visser, M., Jefferies, E., & Lambon Ralph, M. A. (2010). Semantic processing in the anterior temporal lobes: A meta-analysis of the functional neuroimaging literature. Journal of Cognitive Neuroscience, 22(6), 1083–1094. doi: 10.1162/jocn.2009.21309

Wagenmakers, E. J., Ratcliff, R., Gomez, P., & McKoon, G. (2008). A diffusion model account of criterion shifts in the lexical decision task. Journal of Memory and Language, 58(1), 140–159. doi:10.1016/j.jml.2007.04.006

Waters, G. S., & Seidenberg, M. S. (1985). Spelling-sound effects in reading - Time-course and decision criteria. Memory & Cognition, 13(6), 557–572. doi: 10.3758/bf03198326

Weekes, B. S. (1997). Differential effects of number of letters on word and nonword naming latency. Quarterly Journal of Experimental Psychology Section a-Human Experimental Psychology, 50(2), 439–456. doi: 10.1080/713755710

Welbourne, S. R., & Lambon Ralph, M. A. (2005). Exploring the impact of plasticity-related recovery after brain damage in a connectionist model of single-word reading. Cognitive, Affective, & Behavioral Neuroscience, 5(1), 77–92. doi: 10.3758/cabn.5.1.77

Welbourne, S. R., & Lambon Ralph, M. A. (2007). Using parallel distributed processing models to simulate phonological dyslexia: The key role of plasticity-related recovery. Journal of Cognitive Neuroscience, 19(7), 1125–1139. doi: 10.1162/jocn.2007.19.7.1125

Welbourne, S. R., Woollams, A. M., Crisp, J., & Lambon Ralph, M. A. (2011). The role of plasticity-related functional reorganization in the explanation of central dyslexias. Cognitive Neuropsychology, 28(2), 65–108. doi: 10.1080/02643294.2011.621937

Whaley, C. P. (1978). Word—nonword classification time. Journal of Verbal Learning and Verbal Behavior, 17(2), 143–154. doi: 10.1016/S0022-5371(78)90110-X

Woollams, A. M. (2015). For richer or poorer? Imageability effects in semantic dementia patients’ reading aloud. Neuropsychologia, 76, 254–263. doi: 10.1016/j.neuropsychologia.2015.03.023

Woollams, A. M., Lambon Ralph, M. A., Madrid, G., & Patterson, K. E. (2016). Do You Read How I Read? Systematic Individual Differences in Semantic Reliance amongst Normal Readers. Frontiers in Psychology, 7, 1757. doi: 10.3389/fpsyg.2016.01757

Woollams, A. M., Lambon Ralph, M. A., Plaut, D. C., & Patterson, K. (2007). SD-squared: on the association between semantic dementia and surface dyslexia. Psychol Rev, 114(2), 316–339. doi: 10.1037/0033-295x.114.2.316

Woollams, A. M., Silani, G., Okada, K., Patterson, K., & Price, C. J. (2011). Word or word-like? Dissociating orthographic typicality from lexicality in the left occipito-temporal cortex. Journal of Cognitive Neuroscience, 23(4), 992–1002. doi: 10.1162/jocn.2010.21502

Yap, M. J., Balota, D. A., Sibley, D. E., & Ratcliff, R. (2012). Individual differences in visual word recognition: insights from the English Lexicon Project. J Exp Psychol Hum Percept Perform, 38(1), 53–79. doi: 10.1037/a0024177

Yap, M., Pexman, P., Wellsby, M., Hargreaves, I., & Huff, M. (2012). An Abundance of Riches: Cross-Task Comparisons of Semantic Richness Effects in Visual Word Recognition. Frontiers in human neuroscience, 6(72). doi: 10.3389/fnhum.2012.00072

Yap, M. J., Sibley, D. E., Balota, D. A., Ratcliff, R., & Rueckl, J. (2015). Responding to nonwords in the lexical decision task: Insights from the English Lexicon Project. J Exp Psychol Learn Mem Cogn, 41(3), 597–613. doi: 10.1037/xlm0000064

Yap, M. J., Tan, S. E., Pexman, P. M., & Hargreaves, I. S. (2011). Is more always better? Effects of semantic richness on lexical decision, speeded pronunciation, and semantic classification. Psychonomic bulletin & review, 18(4), 742–750. doi: 10.3758/s13423-011-0092-y

Yap, M. J., Tse, C.-S., & Balota, D. A. (2009). Individual differences in the joint effects of semantic priming and word frequency revealed by rt distributional analyses: The role of lexical integrity. Journal of Memory and Language, 61(3), 303–325. doi: 10.1016/j.jml.2009.07.001

Yi, H. A., Moore, P., & Grossman, M. (2007). Reversal of the concreteness effect for verbs in patients with semantic dementia. Neuropsychology, 21(1), 9–19. doi: 10.1037/0894-4105.21.1.9

Zorzi, M., Houghton, G., & Butterworth, B. (1998). Two routes or one in reading aloud? A connectionist dual-process model. Journal of Experimental Psychology: Human Perception and Performance, 24(4), 1131–1161. doi: 10.1037/0096-1523.24.4.1131

